# Behavioral evidence for nested central pattern generator control of *Drosophila* grooming

**DOI:** 10.1101/2020.09.15.298679

**Authors:** Primoz Ravbar, Neil Zhang, Julie H. Simpson

## Abstract

Central pattern generators (CPGs) are neurons or neural circuits that produce periodic output without requiring patterned input. More complex behaviors can be assembled from simpler subroutines, and nested CPGs have been proposed to coordinate their repetitive elements, organizing control over different time-scales. Here, we use behavioral experiments to establish that *Drosophila* grooming may be controlled by nested CPGs. On a short time-scale (5-7 Hz), flies execute periodic leg sweeps and rubs. More surprisingly, transitions between bouts of head cleaning and leg rubbing are also periodic on a longer time-scale (0.3 - 0.6 Hz). We examine grooming at a range of temperatures to show that the frequencies of both oscillations increase – a hallmark of CPG control – and also that the two time-scales increase at the same rate, indicating that the nested CPGs may be linked. This relationship also holds when sensory drive is held constant using optogenetic activation, but the rhythms can decouple in spontaneously grooming flies, showing that alternative control modes are possible. Loss of sensory feedback does not disrupt periodicity but slows the longer time-scale alternation. Nested CPGs simplify generation of complex but repetitive behaviors, and identifying them in *Drosophila* grooming presents an opportunity to map the neural circuits that constitute them.

## Introduction

Animals combine simpler movements into complex routines, forming behaviors with organization across multiple time-scales. For example, a California spiny lobster explores its olfactory environment by waving different segments of its antenna with different frequencies: slow and broad oscillations originating from the base allow the antenna to cover a large space around the animal. The next segment oscillates a little faster, ensuring optimal sampling for local exploration, and the most distal segments, where the sensory organs are located, oscillate fastest, adding fine granularity to the lobster’s olfactory image of the world (Ravbar, field observations). How is a complex behavior assembled from simpler movements in such a harmonious manner?

Central pattern generators (CPGs) are neural circuits that produce rhythmic motor outputs in response to a trigger without requiring ongoing descending drive or patterned sensory inputs (Hooper & Büschges, 2017). CPGs control short stereotypic actions in cat walking, crayfish swimming, locust flight, leech heartbeat, and the stomatogastric and pyloric rhythms of crustaceans (reviewed in Berkowitz, 2019; Grillner, 2006; Marder & Calabrese, 1996; Mulloney & Smarandache, 2010; Selverston, 2010).

However, CPGs may also contribute to control of more complex behaviors. When the movements that compose a behavior repeat, it is inefficient to initiate each step with a separate decision. Automating the sequence by calling its actions in series produces reliable execution. Increasingly complex sequences can be assembled from shorter elements, suggesting hierarchical control. When repetitive subroutines are themselves composed of simpler periodic movements, they may be controlled by nested CPGs, hierarchically organized so that a “high-level” slow CPG controls the behavior on coarse scale and a “lower-level” fast CPG adds the fine structure (Berkowitz, 2019). In other words, the slow CPG controls alternations *between* subroutines and the fast CPG controls alternations *within* these subroutines. Various combinations of coarse and fine oscillators could produce behaviors of arbitrary complexity while still keeping them well-timed, stereotyped, and coherent. Bird song, for example, contains sound syllables executed in sequences. The syllables are short repeating elements, and sequences of syllables make phrases or words that also repeat, creating structure over several time-scales. Ingenious local cooling experiments of specific brain regions cause the whole song to slow down, indicating that it is governed by central pattern generating circuits (Long & Fee, 2008).

Here we show that *Drosophila* grooming behavior contains periodic elements over several time-scales of which we explore two: a fast repeat of individual leg movements (including head sweeps and front leg rubs) and a slow alternation between bouts of head cleaning and front leg rubbing. We demonstrate that both of these repeated elements show evidence of CPG control, and that the two rhythms are usually coordinated, establishing fly grooming as a model system for understanding the circuit architecture of nested CPGs.

## Results

### Two time-scales of grooming are periodic

When flies are covered in dust, they initially groom anterior body parts using their front legs (Seeds et al., 2014). They alternate between bouts of head sweeps, where the legs move synchronously, and bouts of leg rubbing, where the legs move in opposition to each other, scraping the dust off. These movements are shown schematically in **Figure 1A**: the purple and orange arrows indicate synchronous in-phase head sweeps and opposing or out-of-phase leg rubs respectively; the thicker blue arrows show alternation between these two leg coordination modes. Bouts of head cleaning (h) are indicated in purple and front leg rubbing (f) in orange on the ethogram (record of behavior actions over time) shown in **Figure 1B**.

**Figure 1:**
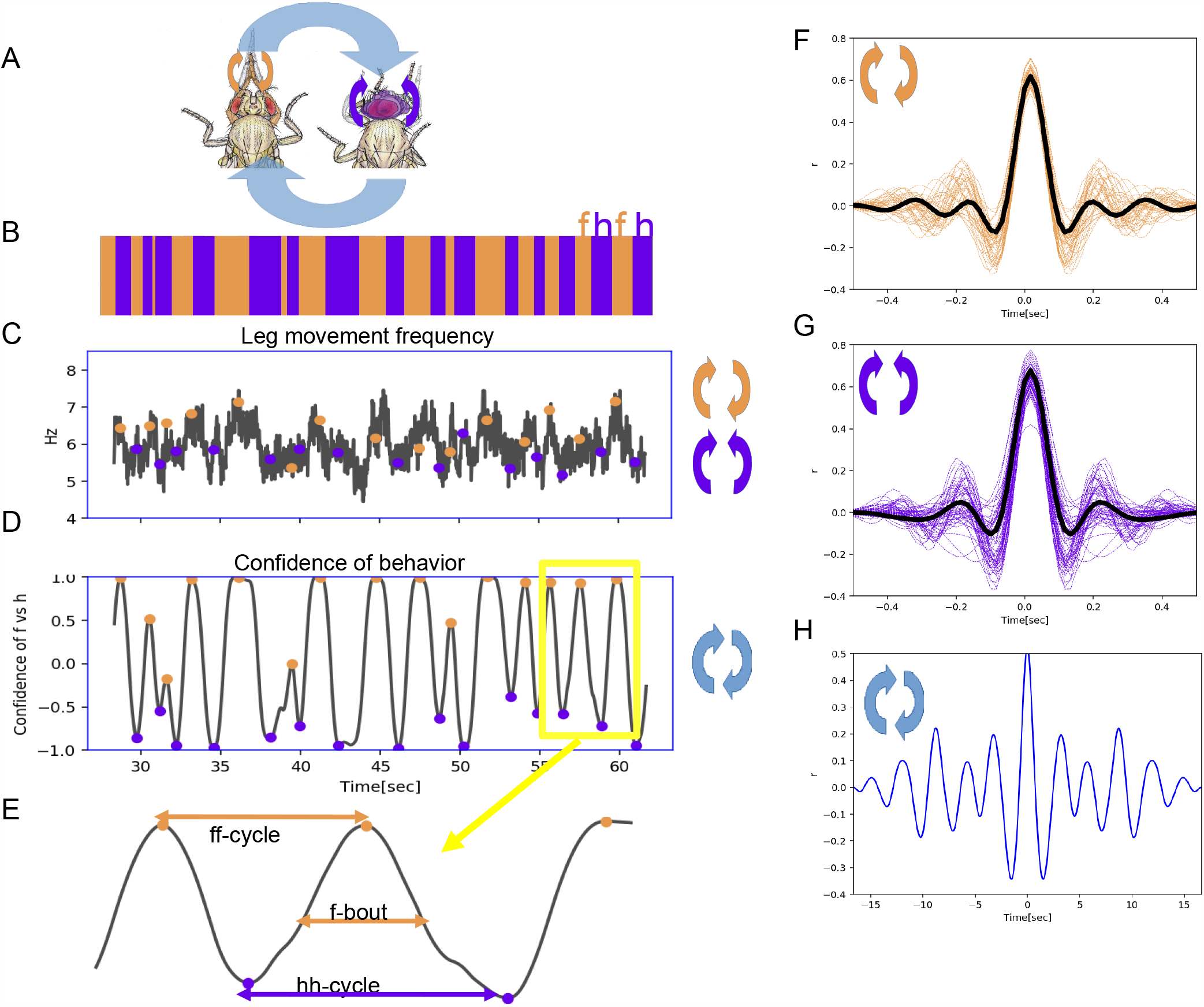
Two time-scales of grooming behavior are periodic. **(A**) Schematic of anterior grooming behavior. The anti-parallel motion of leg rubbing is indicated by the orange arrows and parallel head cleaning movements are indicated by purple arrows (the short time-scale). The blue arrows indicate alternations between the leg rubbing and head cleaning subroutines (the long time-scale). (**B**) Ethogram showing alternations between bouts of front leg rubbing (f) and head cleaning (h) in dust-covered flies recorded at 18°C. (**C**) Example leg sweep and rub frequencies measured in the 30 seconds of anterior grooming behavior shown in the ethogram above. Purple and orange dots indicate front leg rubbing (f) and head cleaning (h) as detected by the Automatic Behavior Recognition System. (**D**) Bouts of front leg rubbing (f) or head cleaning (h) are identified by their probabilities (from the output of the Convolutional Neural Network). When we subtract the probability of h-bouts from that of f-bouts, we obtain the confidence of behavior curve shown here (see Methods). Purple and orange dots indicate maxima and minima of confidence in behavior identification, corresponding to the centers of the f- and h-bouts respectively. (**E**) Enlarged segment taken from (**D)** showing the definitions of ff- and hh-cycles and f-/h-bouts. (**F**) Samples of autocorrelation functions (ACF) computed from over 3 minutes of movies when the fly was engaged in front leg rubbing or head sweeps (**G**). The thick black lines indicate the average of these samples, while thinner purple and orange lines represent each individual ACF contributing to this average.; see Methods for details. (**H**) Autocorrelation function of the alternation of f and h bouts from the example of ff-cycles shown in **D** also reveals periodic signal. **See also Figure 1—figure supplement 1, 2, 3 and 4**

The individual leg sweeps and rubs are stereotyped: these movements are recognizable by human observers or machine vision algorithms (Mathis, 2018; Ravbar, Branson, & Simpson, 2019), and they represent the short time-scale we consider here. We first count individual leg movements from raw videos as they are processed for our Automatic Behavior Recognition System (ABRS) pipeline (see **Methods** and **Figure 1 – figure supplement 1**) and compute their frequencies. In **Figure 1C** we show an example of frequencies of leg sweeps and rubs during the same one-minute period as **Figure 1B**. At 18°C leg rubs and sweeps have a characteristic frequency ~6Hz. This means that each leg movement takes approximately 150msec to complete, which is consistent with our observations using higher resolution video recordings (see below).

Grooming is not a fixed action pattern and flies choose subroutines such as head sweeps or wing cleaning stochastically, but with different levels of prioritization (Seeds et al., 2014). They typically make several head sweeps in a row, followed by repeated leg rubbing movements; we call these chunks of repeated actions *bouts*. We use ABRS (Ravbar et al., 2019) to automatically classify different grooming actions and to identify the time-points of maximum confidence as the centers of bouts (orange and purple circles in **Figure 1D**). Head cleaning bouts and front leg rubbing bouts alternate: we define the time between two head cleaning bouts as an *hh-cycle* while the time between two consecutive front leg rubbing bouts is an *ff-cycle*. An *ff-cycle* will include both leg rubs and head sweeps, which is why we consider the average frequency of both types of leg movements in our analysis. These terms are illustrated in **Figure 1E**, and the cycles are the long time-scale we investigate here.

We demonstrate the periodicity of head sweeps and leg rubs (short time-scale) using autocorrelation analysis (**Figure 1F** and **G)**. Although previous work revealed some syntactic organization at the bout level – both identity and duration of current action influence the identity and duration of the next (Mueller, Ravbar, Simpson, & Carlson, 2019) - we were surprised to find that the alternation of head cleaning and front leg rubbing bouts is also periodic. Autocorrelation analysis of the ff-cycles shown in **Figure 1H** illustrates signal at ~0.45 Hz, corresponding to approximately two seconds between the mid-point of consecutive bouts of front leg rubbing.

To quantify the strength of periodicity, we compare the height of the central peak of the autocorrelation function (ACF) at zero lag to the height of the first prominent neighboring peak (**Figure 1 – figure supplement 2**); we call the ratio of peak heights the Periodicity Index (PI). More periodic movements have high shoulder peaks: a perfect sine wave would have a PI of 1, while in more weakly periodic data, this index would approach 0. The weakest periodic grooming movements we observe have values ~0.2. Non-periodic signals have poorly-defined shoulder peaks and fall below our threshold for peak detection; therefore, the PI is not computed for non-periodic behaviors (see Methods). Using this metric, head sweeps and leg rubbing movements of dusted flies grooming at 18°C have a PI of ~0.33; the alternation between bouts (the long time-scale) has a PI ~0.35. We also compute the amount of periodic behavior as a ratio of periodic to non-periodic behaviors, as shown in **Figure 1 – figure supplement 2**. Using these ratios, we found that overall the amount of periodic behavior for the short time-scale is ~80% and the long time-scale is ~50% at 18°C. For a comprehensive comparison of PI values and the overall amounts of periodic behaviors for the different conditions evaluated throughout this project see **Figure 1 – figure supplement 3**.

We confirm the frequency and periodicity of leg movements by analyzing an independent video dataset analyzed using Deep Lab Cut (DLC), a method developed for tracking of individual body parts (Mathis, et al 2018). **Figure 1 – figure supplement 4** shows the changes of joint angles (resulting from leg movements) over two seconds of grooming behavior: at 18°C, the movement frequencies for the short- and long time-scales are 4.5Hz and 0.22 Hz respectively, similar to our original measurements using ABRS. The PI values are both ~0.34 and the prominence of shoulder peaks is above our threshold for periodic behaviors. The discovery that both short time-scale and long time-scale subroutines within grooming behavior show periodicity suggests the possibility that they may both be controlled by central pattern generating circuits.

### The period lengths of both time-scales contract with increasing temperature

A key feature of central pattern generators is that they oscillate faster at higher temperatures (Deliagina, Orlovsky, & Pavlova, 1983; Tang et al., 2010). To determine whether temperature affects the periodicity of leg sweeps and rubs (short time-scale) or the alternation between bouts of head sweeps and leg rubs (ff-cycles; long time-scale), we recorded the grooming behavior of dust-covered flies at a range of temperatures between 18 and 30°C. Example ethograms from the extreme temperatures are shown in **Figure 2A**, and ethograms from the entire dataset arranged from coolest to warmest temperature of 84 individual flies at seven temperatures recorded for 13 minutes each is displayed in **Figure 2B**.

**Figure 2:**
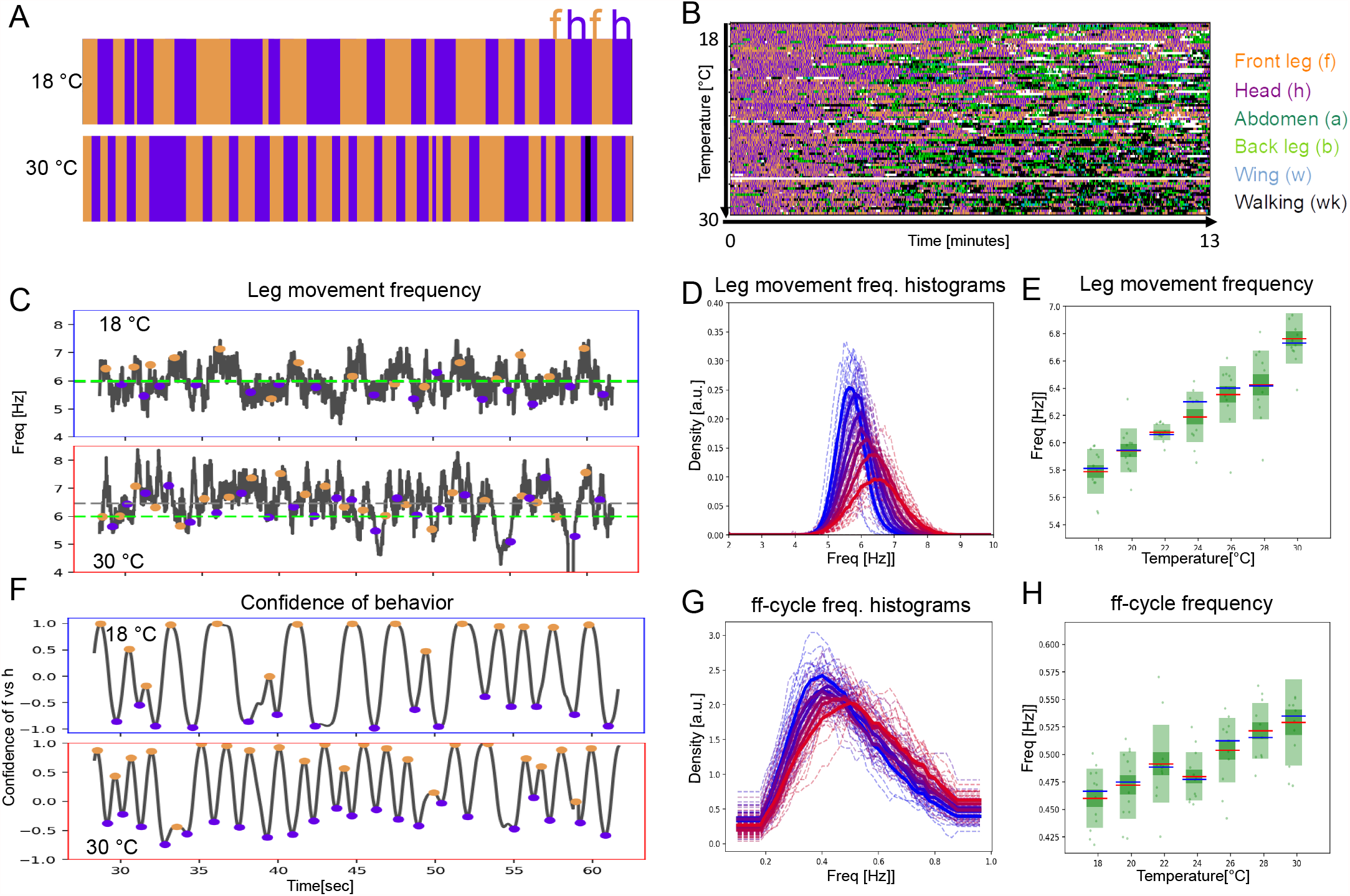
Period lengths of two time-scales contract with increasing temperature in dust-stimulated flies. (**A**) Examples of ethograms recorded at 18°C (top) and 30°C (bottom). (**B**) Ethograms of 84 dust-stimulated flies recorded at different temperature (18 – 30°C). Colors represent behaviors as indicated in the color legend on the right. (**C), (D)**, and (**E**) The frequency of individual leg movements increases with temperature. (**C**) Shows example frequency time series from 30 seconds of grooming at 18° (top, blue outline) and 30°C (bottom, red outline). Gray dashed line = mean of this sample; green dashed line = reference at 6 Hz. (**D**) Histogram of leg movement frequencies, sampled from seven temperatures (18° - 30°C.) Lower temperatures are indicated in blue and higher ones in red. Thin lines – individual histograms; thick lines – average of samples at each temperature. All histograms are computed from the 84 ethograms of dust-stimulated flies in (**B). (E)** Box plots of leg sweep frequencies. Dots show individual fly averages, while the blue bars show the mean frequency, – the red bars mark median and the green shaded areas indicate standard deviation (SD) and error (Blue/red bars in box plots mean/median; shaded areas – SD and SE). (**F**), (**G**) and (**H**) The frequency of ff-cycles also increases with temperature. **(F)** As described in **Figure 1D**, this plot shows the confidence, in samples recorded at 18°C (top) and 30°C (bottom). (**G**) and (**H**): Similar to the panels (**D**) and (**E**) but showing the increase of ff-cycle frequency with temperature computed from the 84 dust-stimulated flies. See **Figure 2-figure supplements 1, 2, 3 and 4**

Temperature increase causes faster individual leg movements (short time-scale) (**Figure 2C-D**) and the frequency shows a linear increase from 5.8Hz to 6.8Hz (R^2^=0.99, p<0.001; **Figure 2E**). This analysis combines sweeps and rubs, but when the different leg movements are considered separately, both show a similar increase with temperature (**Figure 2 – figure supplement 1**).

The period of long time-scale movements is also compressed by temperature. The ff-cycle frequency increases from 0.45Hz at 18°C to 0.53Hz at 30°C, also in a linear manner (R^2^=0.98, p<0.001; **Figure 2F-H**). The frequencies of hh-cycles show a very similar trend (**Figure 2 – figure supplement 2**). Autocorrelations for the long time-scale are more variable at higher temperature, but the alternation remains periodic across the entire range of temperatures (**Figure 1 – figure supplement 3F, Figure 2 – figure supplement 3**). A full spectral analysis of the effect of temperature on the autocorrelation function shows complex structure over several timescales (**Figure 2 – figure supplement 4**).

Increasing temperature shortens the cycle period of both the short time-scale leg sweeps and rubs and the long time-scale alternation between bouts of head cleaning and front leg rubbing. Next, we asked if the two time-scales contract with temperature by the same amount which could suggest a linkage between them.

### Two time-scales contract together with temperature elevation

Several metrics indicate that the short and long time-scales contract at the same rate as temperature increases. We noticed that the number of leg movements within an ff-cycle is fairly consistent (average ~12). **Figure 3A** demonstrates that this average number of leg movements per ff-cycle holds, even as the speed ff-cycles increases with temperature: at 18°C, there are 12 leg movements of 170 msec for a ff-cycle duration of 2.17 seconds, while at 30°C there are 12 leg movements of 148 msec for a ff-cycle duration of 1.89 seconds. Thus, the average number of leg movements per ff-cycle remains the same across all temperatures (**Figure 3B**).

**Figure 3:**
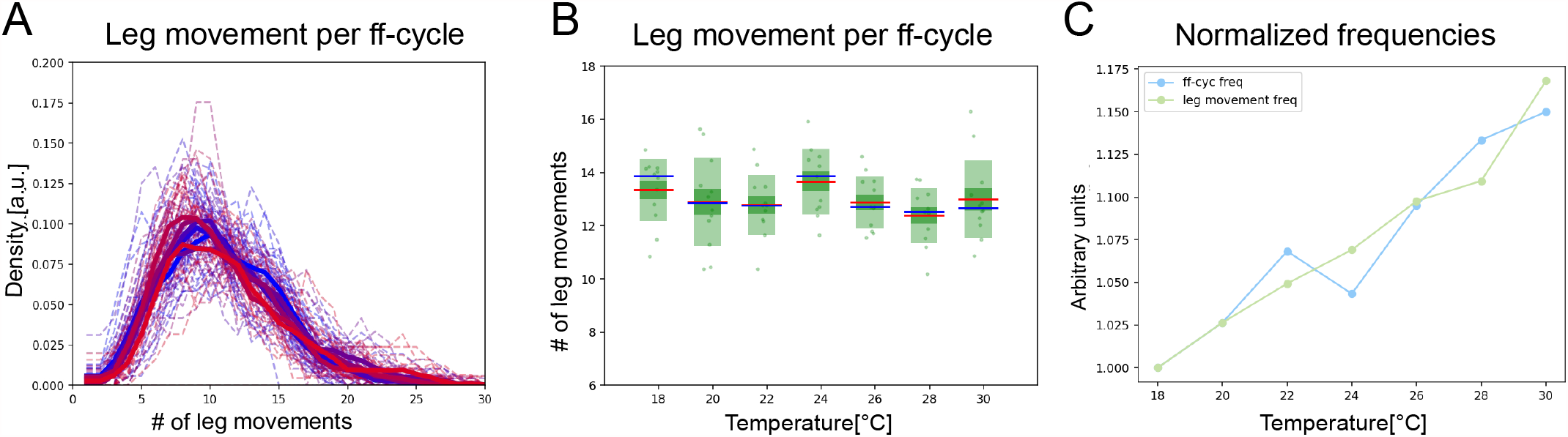
The two time-scales contract together with increasing temperature. (**A**) Histograms of numbers of leg movements per ff-cycle in cooler temperatures (blue) and warmer temperatures (red). (**B**) Box plots of leg movement counts per ff-cycle across the seven temperatures; statistics as in **Figure 2E**. (**C**) The frequency of individual leg movements and bout alternations (ff-cycles) increases roughly linearly with temperature but over different time-scales (msec vs. sec; 7 Hz vs. 0.5 Hz). To see if they increase at the same rate, we compare them in arbitrary units. Frequencies were normalized by dividing each mean value from **Figure 2E** by the lowest value recorded: this produces the rate of change, where 1.0 means no change and values above 1.0 reflect the increased rate. See **Figure 3 – figure supplement 1** for similar effects in hh-cycles.

An alternative way to determine whether the two time-scales scale together with temperature is to plot their contraction *rates*. Normalizing by their minimal frequencies, we can visualize the slope of temperature dependence for each time-scale and the correlation between them is striking (R^2^=0.94, p<0.002; **Figure 3C)**. Both time-scales contract at the same rate, with R^2^ values of 0.96 and 0.93, respectively. Similar trends can be seen for hh-cycles as well (**Figure 3 – figure supplement 1**).

It is possible that temperature will affect both time-scales of grooming behaviors at the same rate just because increased temperature tends to speed up all behaviors through its effect on neural activity, making the apparent coupling between grooming CPGs an epiphenomenon. We consider this unlikely because the two time-scales can be decoupled in spontaneously grooming flies, where they respond differently to temperature, as described below.

### Periodicity and correlation between time-scales persist when sensory stimulation is constant

So far, we have shown that two hallmarks of CPGs – periodicity and temperature-dependent frequency increase – hold for both short time-scale leg sweeps or rubs and long time-scale alternations between leg rubbing and head cleaning subroutines. An additional criterion for determining if a behavior is controlled by a CPG is that rhythmic output does not require rhythmic input. It is challenging to isolate the contribution of sensory input or feedback to the rhythms we observe in grooming. There are mechanosensory bristle neurons that detect dust and induce grooming, and proprioceptive sensors that detect limb position or movement during walking and grooming. If these sensory inputs are rhythmic, they could contribute to both short and long time-scale rhythms.

When flies are covered in dust, their own grooming actions alter sensory input stimuli. The faster their legs sweep, the more quickly dust is removed. Perhaps the coupling between time-scales can be explained because faster leg sweeps result in more dust removal, which reduces sensory drive and thus shortens grooming bouts. In other words, the rhythmic behavioral output could result in similarly rhythmic sensory input - thus not excluding a reflex chain explanation of the observed periodicity (Marder & Bucher, 2001). What happens to the periodicity of the long time-scale grooming rhythms, across temperatures, if the sensory drive is held constant? We predict that if it is indeed CPG-controlled, it should still contract with increasing temperature. We test this using optogenetic activation of all mechanosensory bristle neurons.

We previously demonstrated that this manipulation induces grooming, beginning with the anterior body parts, and alternating between bouts of head cleaning and front leg rubbing (Hampel, McKellar, Simpson, & Seeds, 2017; Zhang, Guo, & Simpson, 2020). Here, we combine optogenetic activation for constant sensory input with changing temperature to show that both individual leg movements and bout-level alternations increase in frequency with temperature and that they do so in a correlated manner. The expression pattern used to activate mechanosensory bristle neurons is shown in **Figure 4 – figure supplement 1**. Representative ethograms at 18° and 30°C show characteristic alternation between bouts of head cleaning and front leg rubbing (**Figure 4A**). The entire behavioral dataset of optogenetically-induced grooming over a range of temperatures is shown in **Figure 4B**.

**Figure 4:**
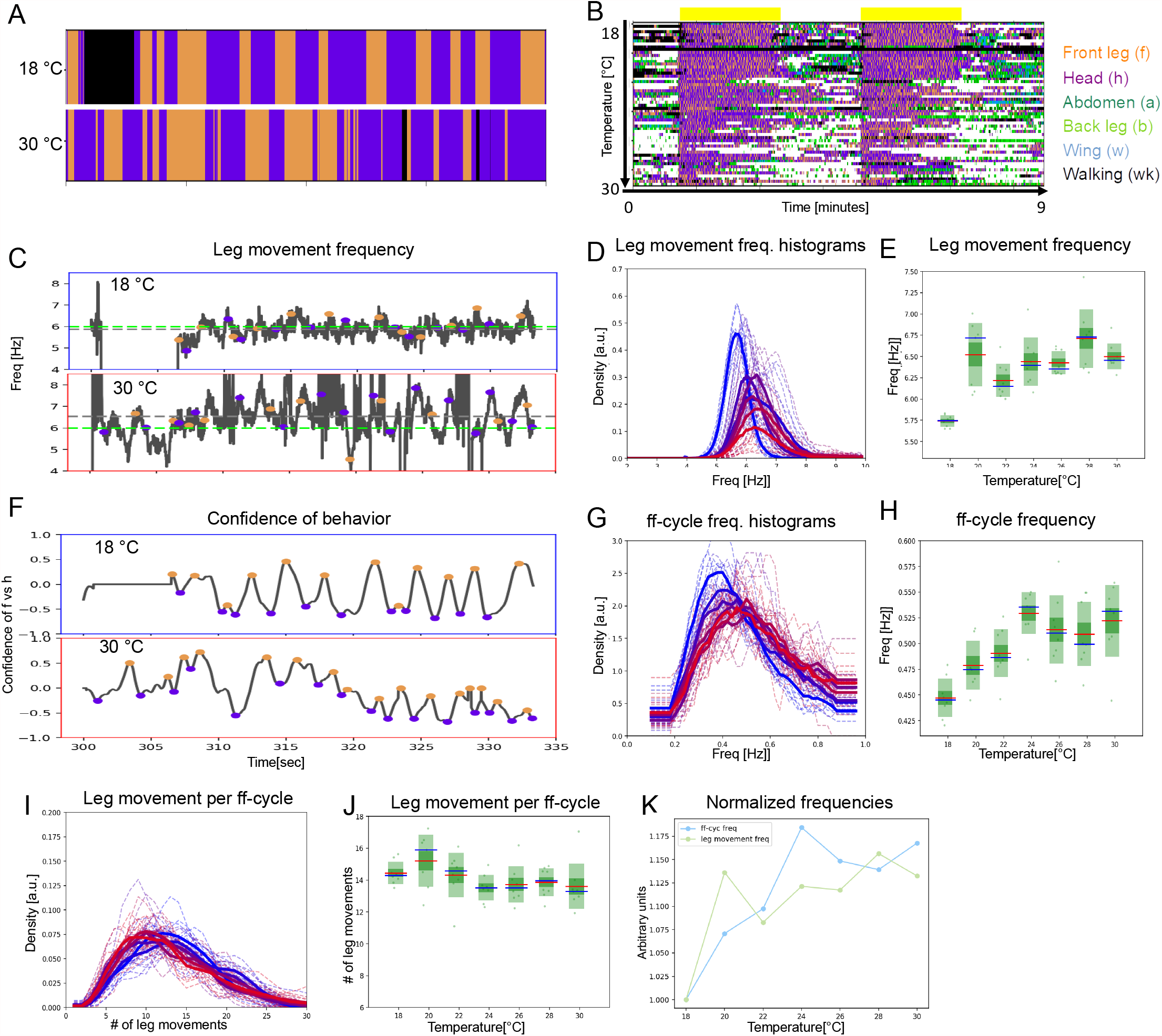
Under constant sensory stimulation, the temperature effect on both time-scales persists. Undusted flies expressing the optogenetic activator *UAS-Chrimson* in mechanosensory bristles were stimulated with red light to induce anterior grooming behavior. Examples ethograms recorded at 18° (top) and 30°C (bottom) are shown in (**A**), while (**B**) shows the whole dataset of ethograms representing 56 flies across the range of temperatures, similar to **Figure 2B**. The yellow bars represent the periods of light activation, lasting 2 minutes each, to optogenetically induce grooming. **(C)** Examples of leg movement frequencies at 18° (top) and 30°C (bottom), (**D**) histograms of mean frequencies at cool (blue) and warm (red) temperatures, and (**E**) box plots of the increase in leg movement frequency with temperature; plots and statistics as described in **Figure 2C, D**, and **E**; compare to short time-scale effects where grooming is induced by dust. (**F, G** and **H**) Long-time scale ff-cycle analysis same as in **Figure 2F, G** and **H. (I)** Histograms and box plots (**J**) of median leg movement counts per ff-cycle across the seven temperatures, quantified as in **Figure 3A** and **B**. (**K**) The rate of temperature-driven increase in frequency is shown by normalization as in **Figure 3C**. See **Figure 4 - figure supplements 1, 2, and 3**.

Uniform sensory input still evokes rhythmic output at both short and long time-scales (**Figure 4 – figure supplement 2**). For example, the periodicity indices at 18°C for short and long time-scales are ~0.35 (**Figure 1 - figure supplement 4E**) and the amount of periodic behavior, considering the long time-scale alterations, is ~60% (**Figure 1 - figure supplement 4H**). This amount of periodic behavior on the long time-scale is less than the ~80% in the dusted flies and the long time-scale rhythm also appears more ragged. We propose that sensory feedback may be needed to stabilize the rhythms but not necessarily to generate them in the first place. The period of the optogenetically-induced rhythms gets shorter with temperature (**Figure 4C-H)**. The frequencies are similar to dust-induced grooming, and the average number of leg movements per ff-cycle is also preserved across the range of temperatures (**Figure 4I-J**). As with dusted flies, the rate of contraction of the two time-scales is correlated (**Figure 4K**), supporting the hypothesis that the short time-scale leg movements and the long time-scale bout alternations are both controlled by CPGs, and that these circuits are yoked together, even under constant sensory stimulation.

Optogenetic activation of mechanosensory neurons does not precisely mimic the physical stimulus of dust itself, and the response of the optogenetically-manipulated flies to temperature reflects this. Both short and long time-scale behaviors occur with somewhat shorter periods at lower temperature than their dust-evoked counterparts, and they stop increasing beyond 26-28° C (compare **Figures 2E, H** and **4E, H**). One possible explanation is that the optogenetic stimulation is “maxing out” the sensory input: it may be driving the fastest leg movements biomechanically possible, or the upper bound of the CPGs frequency range may be reached at a lower temperature. We investigated this by activating the mechanosensory bristle neurons at a constant temperature but with a range of light intensities: the frequencies of movements induced remain constant (**Figure 4 – figure supplement 3**). Even starting from this higher frequency baseline, the two time-scales still increase with temperature at similar rates (**Figure 4K**), indicating that optogenetic activation does not immediately induce maximum movement speeds.

### Periodicity of both time-scales is preserved under reduced sensory feedback

Optogenetic activation may mask acute changes in sensory feedback as the legs contact the body or each other during grooming movements, but removing or silencing sensory neurons is a more direct test of their potential contributions to rhythmicity. To determine if leg rubs and head sweeps remain rhythmic when sensory feedback is reduced, we amputated one front leg between the femur and tibia, similar to Berendes et al, 2016. This eliminates distal proprioceptive feedback from the amputated leg, as well as the usual mechanosensation provided by contact between the legs or the leg and the head during rubs and sweeps. We then employed the DLC software to track the position of the stump and of the intact front leg (**Figure 5 – figure supplement 1**). We found that movements of both the intact leg and the stump remained periodic, with frequencies and periodicity indices similar to those of intact legs in dusted flies. This result further supports that at least the short time-scale is indeed controlled by CPGs.

**Figure 5:**
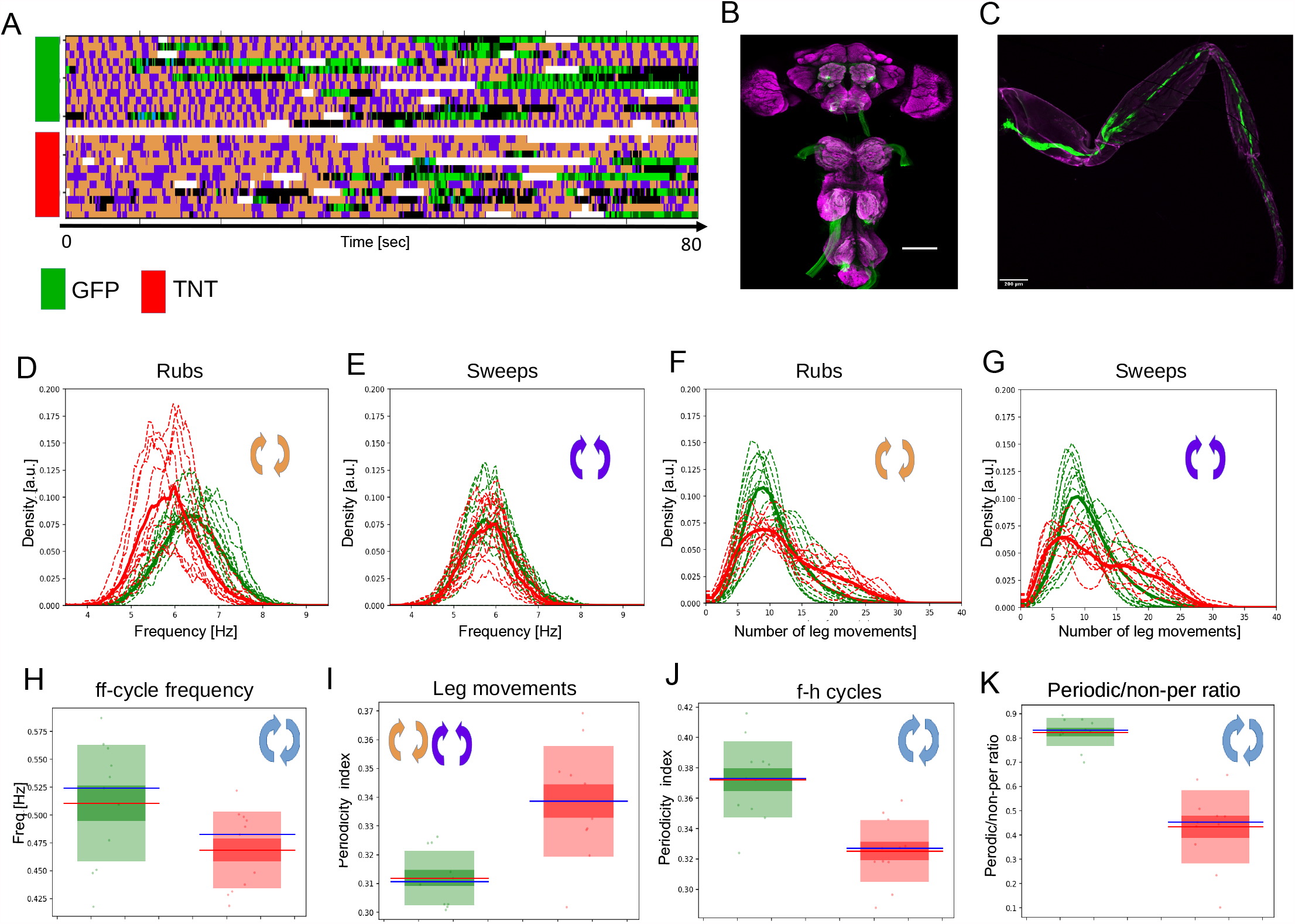
Periodicity of both time-scales is preserved under reduced sensory feedback. (**A**) Ethograms of grooming behavior flies with inhibited proprioception (TNT, red bar) vs the ethograms of the control group (GFP, green bar). (**B**) Expression pattern of the leg mechanosensory neurons driver line used in TNT inhibition experiments. Expression pattern of *540-GAL4, DacRE-flp > 10X-stop-mCD8-GFP* in central nervous system. Magenta: anti-Bruchpilot. Green: anti-GFP. Scale bars, 100 mm. (**C**) Expression pattern of *540-GAL4, DacRE-flp > 10X-stop-mCD8-GFP* in leg sensory neurons. Magenta: cuticle autofluorescence. Green: innate GFP fluorescence. Scale bars, 200 mm. (**D**) and (**E**) Histograms of leg-rub and head-cleaning frequencies, respectively. Green histograms: GFP control flies, red histograms: TNT flies. (**F**) and (**G**) Similar as in **D** and **E** but for the count of rubs and sweeps per ff-cycle. (**H**) Frequencies of the long time-scale (computed from ff-cycles as in **Figure 2G** and **H**) for both groups. (**I**) Box-plots of periodicity indexes for the short time-scale of the control (green) and the experimental (TNT) (red) flies. (**J**) Similar as in **I** but for the long time-scale (f-h oscillations). (**K**) Ratios of periodic vs non-periodic long time-scale oscillations in control and experimental groups. See also **Figure 5 – figure supplement 1** for removal of sensory cues by amputation.

We also attempted to reduce sensory feedback using genetic reagents. We blocked chemical synaptic transmission in leg mechanosensory neurons by expressing tetanus toxin and observed grooming behavior in response to dust. **Figure 5A** shows ethograms of grooming behaviors for both the experimental (TNT) and control (GFP) groups; **Figure 5B-C** shows the neurons that have been genetically inhibited. The frequencies of sweeps and rubs for both groups for the short time-scale are similar (**Figure 5D-E**), but the TNT group showed slower long time-scale oscillations with more leg movements (head sweeps and leg rubs) per ff-cycle than the control group (**Figure 5F-H**). Both groups exhibited periodic behaviors on both the long and short time-scales, but the TNT group was more periodic on short time-scale (**Figure 5I**) and less periodic on the long time-scale (**Figure 5J**). The sensory inhibited flies also performed less overall periodic behaviors on the long time-scale (**Figure 5K**). Taken together, these results suggest that grooming behaviors do not require sensory input for periodicity, but do utilize it for timing, modulation, and perhaps stabilization of the motor output, especially on the longer time-scale.

### Nested CPGs can be decoupled in spontaneously grooming flies

Flies groom robustly in response to dust or optogenetically controlled mechanosensory stimulation, but they also groom spontaneously. The leg movements they perform are recognizable sweeps and rubs, and they occasionally produce alternating bouts of head cleaning and front leg rubbing as well (**Figure 6A, B**). These flies have no experimentally-applied sensory stimuli - only what they generate themselves by contact between their legs and bodies, and the associated proprioceptive feedback - so these motor patterns are most likely to be generated by internal circuits. We analyzed the temperature response of both time-scales of grooming movements in these spontaneously grooming flies.

**Figure 6:**
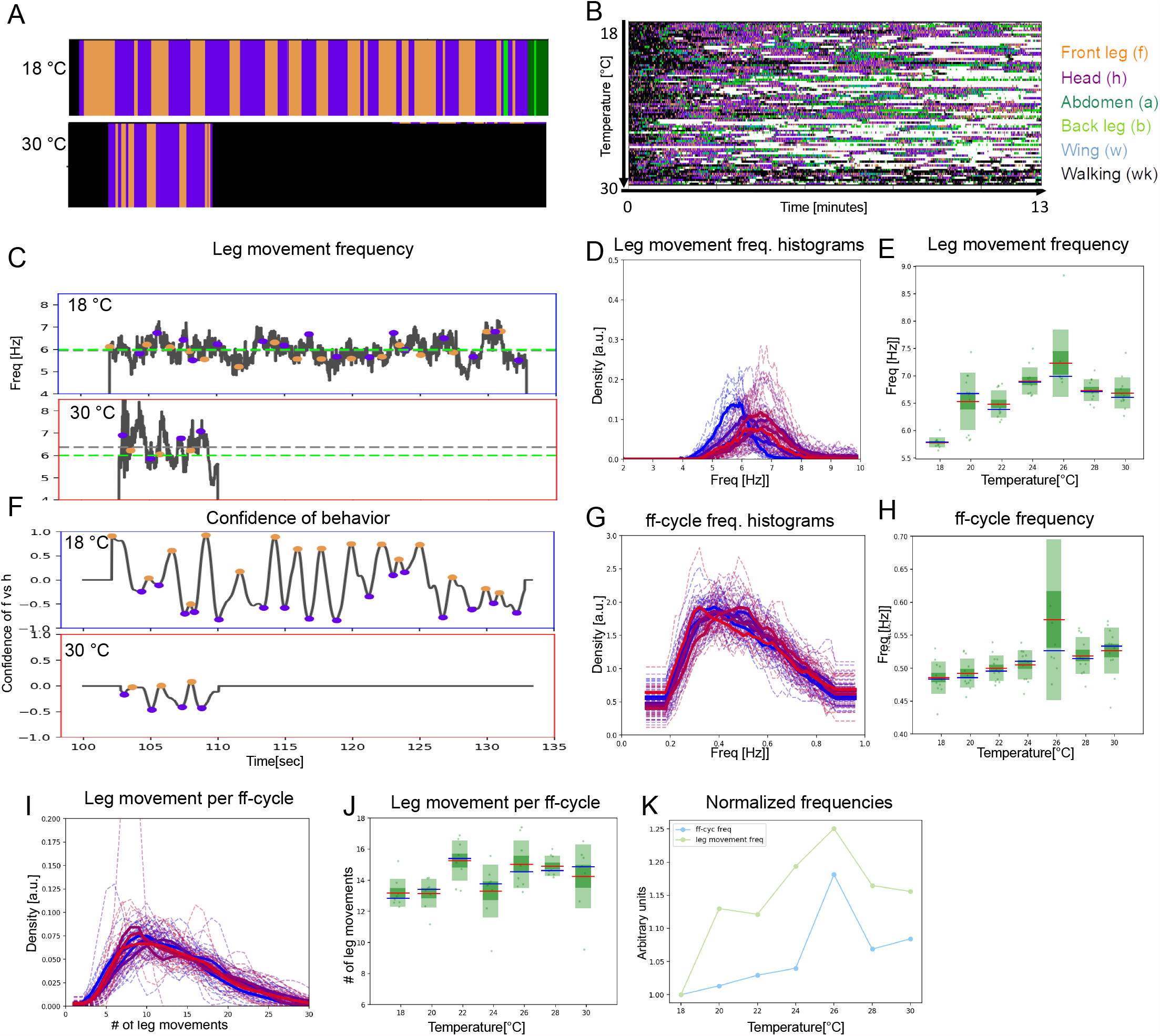
Two time-scales of patterned movements can be decoupled in spontaneous grooming. Spontaneous grooming was recorded in undusted flies at a range of temperatures between 18° and 30°C. Examples are shown in (**A**) and the whole dataset of 80 flies recorded for 13 minutes is shown in (**B**). (**C**) Examples of spontaneous leg movement frequencies at 18° (top) and 30°C (bottom), (**D**) histograms of mean frequencies at cool (blue) and warm (red) temperatures, and (**E**) box plots of the increase in leg movement frequency with temperature; plots and statistics as described in **Figure 2** and **3C, D**, and **E**; compare to short time-scale effects where grooming is induced by dust. (**F, G** and **H**) Long-time scale ff-cycle analysis comparable to **Figure 2** and **3F, G** and **H. (I)** Histograms and box plots (**J**) of median movement counts per ff-cycle across the seven temperatures, quantified as in **Figure 3A** and **B**. (**K**) The rate of temperature-driven increase in frequency is shown by normalization as in **Figure 3C**. See also **Figure 6 – figure supplements 1, 2, and 3** for evidence of periodicity and response to temperature.

Although these flies groom less than dusted or optogenetically activated flies, they show characteristic sweep and rub frequencies that increase with temperature, albeit with higher variance (5.5Hz to 7.2Hz; **Figure 6C-E**). The ff-cycles are rarer, and when they occur, the temperature-dependent contraction is much less pronounced (**Figure 6F-H**). Although the number of leg movements per ff-cycle is similar to stimulated flies (~14; with higher variation across the temperature range, **Figure 6I, J**), the correlated temperature-dependent contraction of the short and long time-scales observed in the dusted and optogenetically activated flies is *not* seen in the spontaneously grooming ones (compare **Figure 6K** to **Figures 3K** and **4K**). The frequency of leg rubs and sweeps increases with temperature at a greater rate than ff and hh-cycles (**Figure 6K**), suggesting that the pattern-generating circuits that control the two time-scales of movements can be dissociated in spontaneous grooming.

Perhaps some aspects of grooming, namely the long time-scale alternations between bouts of head cleaning or front leg rubbing, can either be driven rhythmically by a high-level CPG (as they seem to be in stimulated flies) or initiated individually in a less patterned or CPG-independent way, in spontaneously grooming flies. To further explore this possibility, we separated the periodic long time-scale oscillations from the non-periodic ones using a threshold of ACF shoulder peak prominence (see **Methods** and **Figure 1 – figure supplement 2**). We hypothesized that if the periodic behaviors are CPG-driven and the non-periodic behaviors are not, then the periodic behaviors should scale with temperature more strongly than non-periodic ones. We found that indeed, while increasing temperature does increase the speed of *all* movements (the bouts, and the intervals between the bouts, get shorter), the frequency of periodic long time-scale oscillations correlate with temperature more significantly than that of the non-periodic f-h alternations (R^2^ = 0.83, p=0.02 vs. R^2^ = 0.7, p= 0.07 respectively), as shown in **Figure 6 – figure supplement 1C** and **D**. We also found that periodic grooming behaviors show a more significant correlation between the short and the long time-scale than the non-periodic behaviors (R^2^ = 0.82, p=0.03 vs. R^2^ = 0.67, p= 0.09 respectively) as shown in (**Figure 6 – figure supplement 1E** and **F**). Taken together these results suggest that grooming has several control architectures with flexible linkage between CPG levels.

## Discussion

Temperature manipulations have been instrumental in identifying behaviors controlled by central pattern generators and for locating where those circuits reside. Robust rhythms are retained across a range of temperatures in the crustacean stomatogastric ganglia, where the relative timing of events is preserved even as the sequence itself changes speeds (Alonso & Marder, 2020; Rinberg et al., 2013; Tang, Taylor, Rinberg, & Marder, 2012). Local cooling of the cat spinal cord demonstrated which segments contain locomotion control circuits (Deliagina, Orlovsky, & Pavlova, 1983), and local warming showed that cricket chirping is governed by thoracic rather than brain ganglia (Pires & Hoy, 1992a, 1992b). The role of CPGs in bird song, and the importance of the HVC brain region for sequence timing, was shown because local cooling expands the entire song without changing the relative durations of the syllables and sub-syllabic components (Armstrong & Abarbanel, 2016; Long & Fee, 2008). In frogs (*X. laevis*) both local cooling and ambient temperature affect the frequencies of vocalization patterns (Yamaguchi et al, 2008). Here, we follow this tradition to examine fly grooming behavior at different temperatures, demonstrating that both its short and long time-scale components show evidence of CPG control.

Although here we change the temperature of the whole fly, our study opens the way to use anatomical and genetic tools to map underlying neural circuits in future experiments.

Fly grooming is an innate motor sequence with both repetition and flexibility. Behavior analysis has shown organization over several time-scales, from single stereotyped leg movements and alternations between bouts of repeated actions targeting specific body parts (Mueller, Ravbar, Simpson, & Carlson, 2019), to a gradual and probabilistic progression from anterior toward posterior grooming (Seeds et al., 2014). Here we investigate what aspects of fly grooming are periodic, demonstrating that the short time-scale leg sweeps and rubs repeat at characteristic frequencies, in the absence of patterned sensory input, and in a temperature-dependent manner (**Figure 2D-E)**, consistent with control by central pattern generators. Since the neural circuits that constitute the CPGs controlling these periodic leg movements are likely to overlap with those proposed to coordinate walking (Bidaye, Bockemuhl, & Buschges, 2018), this was not unexpected.

But our more surprising discovery was that the alternation between bouts of head cleaning and front leg rubbing is also periodic (**Figure 1 – figure supplement 4A**). While this oscillation is more variable, it too, is independent of patterned sensory input, and increases in frequency with temperature (**Figure 2G-H, Figure 4G-H)** – suggesting that there may be an additional CPG operating at this longer time-scale. This high-level, overarching, CPG can control the alternations between leg rubs and sweeps that are themselves governed by faster, low-level CPGs. The idea that multiple CPGs coordinate movements is not new: CPGs may control each leg joint, regulating the interaction between flexor and extensor muscles, governing the way coxa-trochanter and femur-tibia joints are coordinated to produce forward or backward walking, or mediating interactions among limbs (Feng et al., 2020; Mantziaris, Bockemuhl, & Buschges, 2020). In the vertebrate spinal cord, inhibitory and excitatory commissural neurons can cause the CPGs controlling the legs to synchronize for a hopping gait or operate out-of-phase for walking (Kiehn, 2016). The concept of nested CPGs has recently been extended to explain flexible coordination of behaviors ranging from fish swimming to bird song (Berkowitz, 2019). We map fly grooming behavior into this nested CPG framework as shown in **Figure 7**.

**Figure 7:**
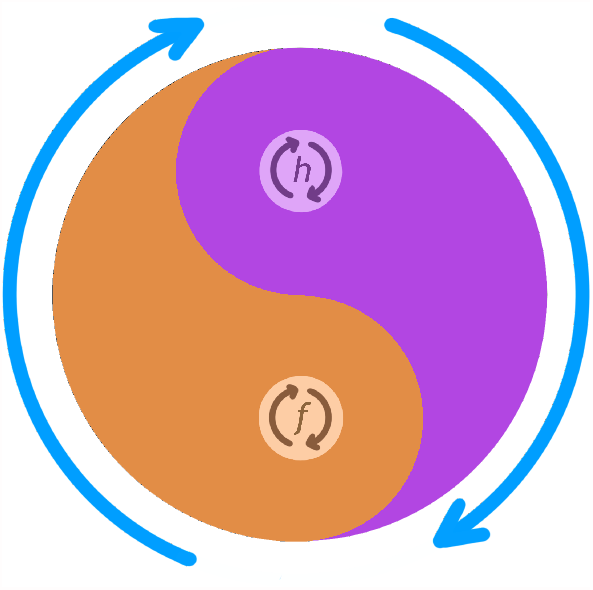
Conceptual schematic of nested CPG controlling anterior grooming behavior. This diagram illustrates how the lower level CPG modes controlling individual leg rubbing (*f*) and head cleaning (*h*) oscillations (small orange and purple arrows) can be coordinated by an over-arching, high-level, CPG controlling the alternation between the front leg rubbing and head cleaning bouts (large blue arrows). The probability of *f* vs *h* is indicated by the thickness of their corresponding colors (orange vs purple) at each angle of the circle. For example at 5 o’clock the probability of *f* is higher than that of *h* and at 11 o’clock probability of *h* is higher than that of *f*, etc.

In a recent publication, nested CPGs similar to the one we are proposing here have been described in the crab stomatogastric ganglion, where fast pyloric and slow gastric mill rhythms are coupled and compensate for temperature changes (Powell, Haddad, Gorur-Shandilya, & Marder, 2020). The combination of two rhythmic behaviors raises the possibility that the governing circuits interact, like meshed gears of different sizes, to simplify control of a repetitive behavior on multiple time-scales. Just as the relative number of rotations between different gears is the same regardless of the absolute speed of the mechanism, so the phase of different components of CPGs is conserved across a range of temperatures (Marder & Bucher, 2001). In the case of fly grooming, this relationship between fast and slow components of the system may be reflected in the constant number of leg movements within each grooming bout across a range of temperatures (**Figure 3**). The fast and slow components of the nested CPG can be coupled and decoupled sporadically over the period of behavior execution which can result in alternations between periodic and non-periodic behaviors on the long time-scale (**Figure 6 – figure supplement 1, Figure 6 – figure supplement 2B**). The neural circuit implementation of nested CPGs, and the “clutch” mechanisms that engage and disengage their connections, remains to be determined.

Dust-induced grooming can be routine, rhythmic, repetitive, and governed by central pattern generators, but spontaneous grooming may have more varied control architecture. Fast walking insects may be more likely to engage CPGs, while slower ones may rely more on sensory feedback and reflex chains (Mantziaris, Bockemuhl, & Buschges, 2020). Just as you can walk a straight path without thinking about it or you can carefully place each foot on icy terrain, flies may have alternative ways to produce grooming movement sequences. Disturbing a single bristle elicits a single directed leg sweep (Kays, Cvetkovska, & Chen, 2014; Vandervorst & Ghysen, 1980), while covering the fly in dust evokes an entire grooming program with bouts of several leg sweeps and rubs, alternation between the bouts, and slow gradual anterior-to-posterior progression (Phillis et al., 1993; Seeds et al., 2014). Our observation that spontaneous grooming shows less periodicity at the long time-scale (**Figure 1 – figure supplement 4G**) can be interpreted as effective decoupling of the two levels of the nested CPG in those flies when they engage in more sporadic and shorter episodes of grooming. This proposal is further supported by observation that the temperature effect is stronger in spontaneously grooming flies when we only consider periodic long time-scale behaviors (**Figure 6 – figure supplement 1**).

In both optogenetically stimulated flies and in flies where sensory feedback was inhibited, we observe less periodicity of the long time-scale than in the dust-stimulated flies (**Figure 1 – figure supplement 2H** and **Figure 5K**). We speculate that sensory feedback is needed to stabilize and modulate the rhythmic behavior, even though it is not necessary to produce it. For example, a fly could produce in-phase rhythms to provide the general control of head-cleaning movements but the sensory feedback from various parts of the head could adjust these movements to optimize the efficiency of cleaning of those parts. The concept of sensory feedback playing a role in stabilizing and adjusting rhythmic behaviors has been explored in robotics (Wang, 2014, Sartoretti, G., 2018) and in walking mice (Mayer W. P., 2018, Santuz, A., 2019), as well as in computational modeling of CPGs (Yu, Z., 2021).

Nested CPGs can be applied to generate arbitrarily complex behaviors that are also periodic and well-coordinated. Many different time-scales of simple periodic movements can be combined into a more complex periodic behaviors the way different frequencies of sine waves can be combined to create arbitrarily complex shapes in Inverse Fourier Transformation. If CPGs can operate across various time-scales in this fashion then all that is needed to create arbitrarily complex yet highly reproducible behavior is adjusting the weight of each contributing level of a multi-level nested CPG.

The specific neural circuits that constitute CPGs have been challenging to identify in most preparations and even the best characterized would benefit from more comparators. The electrophysiological recordings that have been so critical in CPG analysis in other preparations are possible but challenging in *Drosophila*, but the real strength of the system is the ability to demonstrate that specific neurons have a causal connection to rhythm generation. New anatomical resources to map neural circuits, especially the complete electron microscopy dataset covering the ventral nerve cord (Phelps et al., 2021), will enable identification of the pre-motor neurons most likely to participate in CPGs and the commissural connections that may mediate among them. Functional imaging of neural activity in dissected, fictive preparations (Pulver et al., 2015) and even in behaving flies (Chen et al., 2018) is another promising approach to identify neurons with rhythmic activity that many constitute parts of the central pattern generators. The genetically encoded calcium indicators and new voltage sensors have fast enough onset and offset kinetics to capture rhythmicity on the expected time-scales (Simpson & Looger, 2018). For example, whole nervous system monitoring of neural activity was recently described in *C. elegans* to identify hierarchical and dynamic control of nested locomotion patterns (Kaplan 2020).

Rhythmic activity and circuit connectivity may suggest candidate neurons; genetic tools to target specific neurons (Jenett et al., 2012) and optogenetic methods to impose altered activity patterns (Klapoetke et al., 2014) presents a way to causally connect specific neurons to control of rhythmic behaviors. The behavioral evidence presented here suggests that a two-level nested CPGs can control aspects of fly grooming over at least two time-scales. Identifying what circuits constitute these central pattern generators and mapping the neurons that coordinate their interactions is feasible, now that we know we ought to be looking for them.

## Acknowledgements

We thank members of the Simpson, Louis, and Kim labs for feedback, and especially Li Guo and Dr. Josh Mueller for excellent suggestions. We thank Carla Ladd for creative input with figure design and Aleks Labuda for instrument design for high resolution video recording. This research was supported by NIH-R01NS110866 and HHMI Janelia transition funding.

## Methods

### Fly stocks and husbandry

*Drosophila melanogaster* were reared on common cornmeal food in a 25°C incubator on a 12 hr light/dark cycle. 3-5 days *CantonS* males were used for dusting and spontaneous grooming experiments. For optogenetic experiments, larvae were raised on normal food. After eclosion, 1-day old adults were transferred into food containing 0.4 mM all-trans-retinal and reared in the dark for another two days. *R74C07-GAL4* (Bloomington Stock Center 39847), *20XUAS-IVS-CsChrimson*.*mVenus* (Bloomington Stock Center 55134), *5-40-GAL4* (Hughes et al 2007), *DacRE-FLP* (Mendes et al 2013), *UAS>stop>TNT* (Stockinger, et al. 2005) and *10XUAS>stop>myr-GFP* (Bloomington Stock Center 55811) were used in the paper. 10-day old males were used in TNT inhibition experiments. 3 to 5-day old males were used in other experiments.

### Behavior experiments with temperature control

Behavior videos were recorded inside New Brunswick I2400 incubator shaker or DigiTherm DT2-MP-47L heating/cooling incubator. Experiments were performed every 2°C between 18°C and 30°C. Temperature was monitored by a Govee H5072 Bluetooth thermometer. For dusting experiments, the room temperature was also adjusted to the target temperature to make sure flies stay at the same temperature during fly dusting. Each fly was tested once in one condition. Three types of chambers were used in fly dusting assay: dusting chamber (24 well corning tissue culture plate #3524), transfer chamber and recording chamber. Flies were anesthetized on ice and transferred to the middle four wells of the transfer chamber. Transfer chamber with flies was then kept in the incubator set to the target temperature for 15 min. For fly dusting, around 5 mg Reactive Yellow 86 dust was added into each of the 4 middle wells of the dusting chamber. Transfer chamber was aligned with the dusting chamber. Flies were tapped into the dusting chamber and shaken 10 times. After dusting, flies and dust were transferred back into the transfer chamber. Transfer chamber was banged against an empty pipette tip box to remove extra dust. Dusted flies were then immediately tapped into the recording chamber for video recording. The whole dusting process was performed in a Misonix WS-6 downflow hood.

For optogenetics and spontaneous grooming experiments, ice-anesthetized flies were put into the recording chamber directly. Recording chamber with flies was then kept in the incubator set to the target temperature for 15 min before experiment. In spontaneous grooming experiments, flies were shaken for 5 times after the 15 min incubation. They were then rested in the incubator for another 3 minutes before recording.

60 Hz videos were recorded for 9 min in optogenetics experiments, 13 min in dusting and spontaneous grooming experiments with a FLIR Blackfly S USB 3 camera. Each video is recorded from the top and it captures flies in four chambers. Infrared backlight was used for all experiments. Custom-made LED panels (LXM2-PD01-0050, 625nm) were used for optogenetic activation from below. 20 Hz 20% light duty cycle was used in optogenetics experiments. Red light intensity was adjusted to 0.85 mW/cm^2^. Before the video analysis the frames were split into four quadrants corresponding to the four chambers.

For all experiments described above, 8-12 flies were used per temperature or per experimental group (for the TNT inhibition experiment). Occasionally a recording chamber was empty or the fly was visibly damaged or dead. Those chambers were excluded from further analysis. These numbers were sufficient to extract the quantities we use in this work such as ff-cycle periods.

### High-resolution videos recording

For high-resolution videos used for leg tracking, flies were put in a 10mm diameter quartz chamber, and 100hz videos were recorded from below. A FLDR-i132LA3 red ring light (626nm) was used for optogenetics activation. For leg amputation experiments, one front leg was amputated at the middle of the femur. Flies were recovered for 3 days before the experiments. The dusting procedure is the same as what described above.

### Leg movements counting from video

We did not track individual legs or joints so the frequencies of leg movements during grooming were estimated from frame-to-frame changes in pixel values. When flies are walking or engaged in other non-grooming behaviors, such measurements would not be very useful. However, during grooming flies are standing in one place and the most intense pixel value differences roughly correspond to those created by leg movements (rather than whole body translations).

Regions of interest were cropped around the animals to produce 80×80 arrays of pixels (**F**). The **F** arrays were reshaped into 1600 columns and 32 rows, corresponding to 32 consecutive frames (~0.5 sec window at 60Hz), were stacked together to obtain 32×1600 array of pixel values (**W**).

The derivatives of **W** across time were computed using Numpy.gradient() function to obtain matrix of frame-to-frame differences **D**. The matrix **D** was smoothed using Gaussian filter (`scipy.ndimage.filters.gaussian_filter`). Sigma = [2, 5]. This was done to denoise the signal - to remove non-biological high frequency changes (temporal smoothing) and to reduce spatial resolution (so that more neighboring pixels capture the same signal). All the values of **D** where cumulative change of pixel values was less than threshold (θ = 5.0) were converted to zeros to retain only those pixels where significant movement occurred. An example of **D** matrix is shown in **Figure 1 – figure supplement 1**

Notice in the figure the several non-zero columns (corresponding to pixels) that appear sinusoidal. These columns contain the signals that have been smoothed to remove the high frequency noise and are prominent enough to “survive” the thresholding. Therefore we assume that the movements represented in these columns correspond to relatively large, periodic actions that occur while the animal is standing (otherwise the sinusoidal signals would not stay in one column over the time-window). Also, only the frames where grooming behavior was detected by the ABRS were considered for leg movement counting. This assumption leads us to conclude that the periodic signals are caused by periodic leg movements during grooming. So the counting the peaks across the columns of matrix **D** would then result in the number of times a leg crossed a particular pixel. The peaks were found applying the `scipy.signal.find_peaks()` function across the rows on the matrix **D**. The peaks were then counted for each column in D and the median of all the counts was finally taken to be the number of leg sweeps or rubs that occurred in that time-window (.5 sec). Note that at the frame rate of 60 Hz we should be able to detect frequencies of periodic signals below 30 Hz and the highest frequency of leg movements we observed was below 7 Hz.

The number of leg movements per second was used as the leg movement frequency.

### Automatic behavior recognition from videos

Probabilities and ethograms of grooming behaviors were extracted from raw videos using Automatic Behavior Recognition System (ABRS). For detailed description see (Ravbar et al., 2019) and for the most updated version see ABRS (https://github.com/AutomaticBehaviorRecognitionSystem/ABRS)

Briefly, ABRS pre-processes video data by compressing it into spatio-temporal features in the form of 3-channel space-time images (3CST images, shape=80 × 80 × 3) where the first channel [0] contains raw video frame pixel values [80×80], the second channel [1] contains spectral features of pixel value dynamics over a time-window of 16 frames, and the third channel [2] contains frame-to-frame difference (frame subtraction). The 3CST images are classified into 7 behavioral categories (front leg rubbing, head cleaning, back leg rubbing, abdominal cleaning, wing cleaning and whole-body movements (walking)) by ConvNet (Covolutional Neural Network - CNN) implemented in Tensor Flow, (in Python) (https://www.tensorflow.org) using a model trained with diverse set of videos of fly grooming behaviors. In the final output layer of the CNN are the probabilities of the grooming behaviors. In this work we focus entirely on the probabilities of front leg rubbing and head cleaning behaviors: P(f) and P(h).

### Behavioral confidence

The long time-scale oscillations are quantified as probabilities of behaviors obtained from the ABRS CNN described above. Behavioral confidence is computed as:

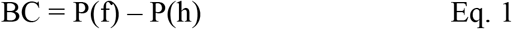

The BC signal is smoothed two consecutive times with time-window of 31 frames (0.5 sec) using the `scipy.signal.savgol_filter` function to remove high frequency noise.

### Autocorrelation and periodicity analysis

Autocorrelation functions (ACFs) were computed from a 0.5 sec time-window for individual leg movements (leg rubs and sweeps). The signal for autocorrelation was extracted from raw movies as follows: 1) Matrix **D** was obtained as described above (Leg movements counting from video); 2) ACFs were computed for every column of **D** (an example of the ACFs and the matrix **D** are shown in **Figure 1 – figure supplement 1**); and 3) All the ACFs were averaged to obtain the mean ACF for that time-window. The mean ACFs were stacked to obtain the ACF array with dimensions 60 × F, where F is the number of frames in the movie. The AFCs were computed using `scipy.signal.correlate` function from SciPy library.

Autocorrelation functions (ACFs) for the long time-scale (the oscillations between bouts of leg rubbing and head cleaning) were computed from the behavioral confidence (BC) time-traces described in Eq.1, in the time-window of 999 frames (16.6 sec), also using the `scipy.signal.correlate` function.

To quantify the periodicity of each time-scale we computed the *Periodicity Index* from the ACFs, defined as the ratio between the height of the central peak of an ACF and the nearest prominent peak (**Figure 1 – figure supplement 2**). The peaks were detected by `scipy.signal.find_peaks` function and the threshold for “prominence” was set at 0.2. We used the same method and parameter settings for both time-scales to compute the Periodicity Index.

We managed to separate periodic behaviors from the non-periodic by using the `scipy.signal.find_peaks` function. If the first neighbor of the central peak of the ACF fell below the 0.2 “prominence” threshold we classified the pertaining behavior as non-periodic and as periodic otherwise. This is shown in **Figure 1 – figure supplement 2B**.

### Frequency computations for the long time-scales

Behavioral confidence (BC) traces were used to compute the frequencies of the long time-scale.

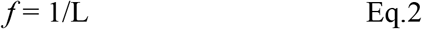

In Eq. 2 the L is a vector containing the lengths of periods measured from the BC traces. The periods were measured either between two peaks of the BC traces (ff-cycles) or between two valleys (hh-cycles) (also see **Figure 1E** for illustration). This produced a vector of frequencies of dimensions 1xF, where F is the number of frames in a movie, corresponding to one fly. The mean and median frequencies for each fly were computed as means/medians of vector *f*.

### High resolution behavior analysis

To confirm our observations of grooming behavior, its periodicity and the effect of temperature, we used an independent method for limb tracking, the Deep Lab Cut (DLC) (Mathis, et al 2018) on a small sample of high-resolution videos described in the previous section. The DLC allowed us to label (virtually) three body parts on each front leg and a reference point (see **Figure 5 – figure supplement 1 A** showing the labels on the intact front leg). We could then track these labels across time. For each leg we computed the angle between Tibia and Femur and tracked the derivatives of these angles across time examples are shown in **Figure 1 – figure supplement 4A**. On these time-traces we applied the auto-correlation analysis and FFT to compute ACFs and their spectra. (For the amputated fly we used the relative positions of the joints and the stump instead of the angles.) We computed the Periodicity Index from the ACFs.

In these analyses, we did not have access to the Behavioral Confidence (see the section above) so, in order to examine the periodicity and frequencies of the long time-scale, we computed the equivalent to behavioral confidence as follows: the distance between tarsal segments was used as the “confidence of head cleaning” and the angle between Tibia and Femur was used as the “confidence of front leg rubbing”. We obtained the behavior confidence signal by subtracting the latter from the former. We applied autocorrelation and FFT analysis this “behavior confidence” to estimate the Periodicity Index and the frequency of the long time-scale. Due to cumbersome nature of these analysis and the lack of behavioral recognition we performed them only on three different movies from the dust-stimulated flies (**Figure 1 – figure supplement 4D-F**).

### Handling of outliers and missing data points

We found and did not eliminate an outlier in optogenetically stimulated flies (leg movement frequencies at 20°C). In cases where there were missing data points (no fly was in the recording chamber), those were not counted in the statistics.

---

**Figure 1—figure supplement 1:**
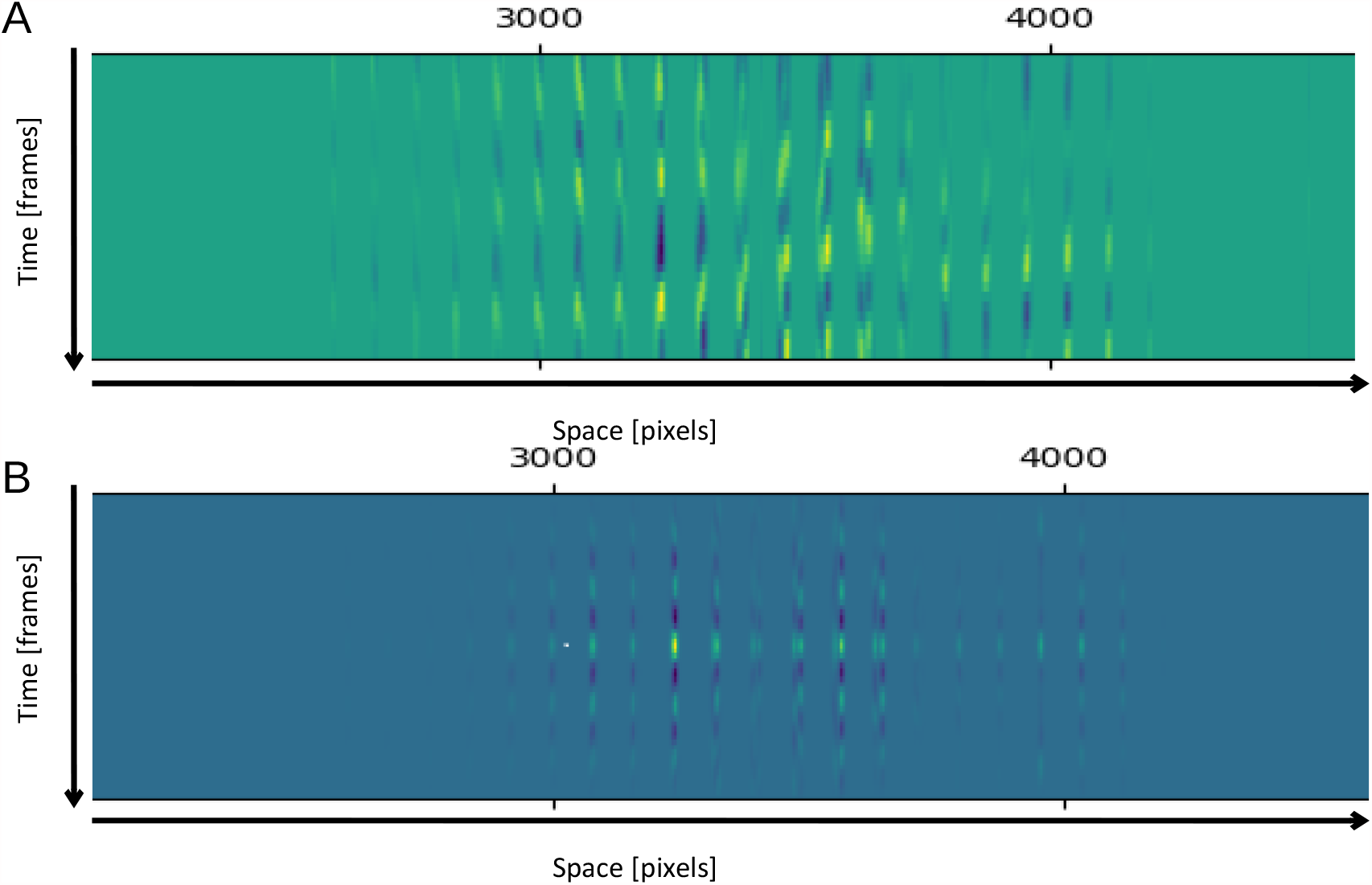
Methods of leg movements (sweeps or rubs) counting from video. (**A**) Smoothed derivatives from matrix ***D*** described in Methods. (**B**) Auto-correlation functions (ACFs) computed from the matrix ***D***.

**Figure 1—figure supplement 2:**
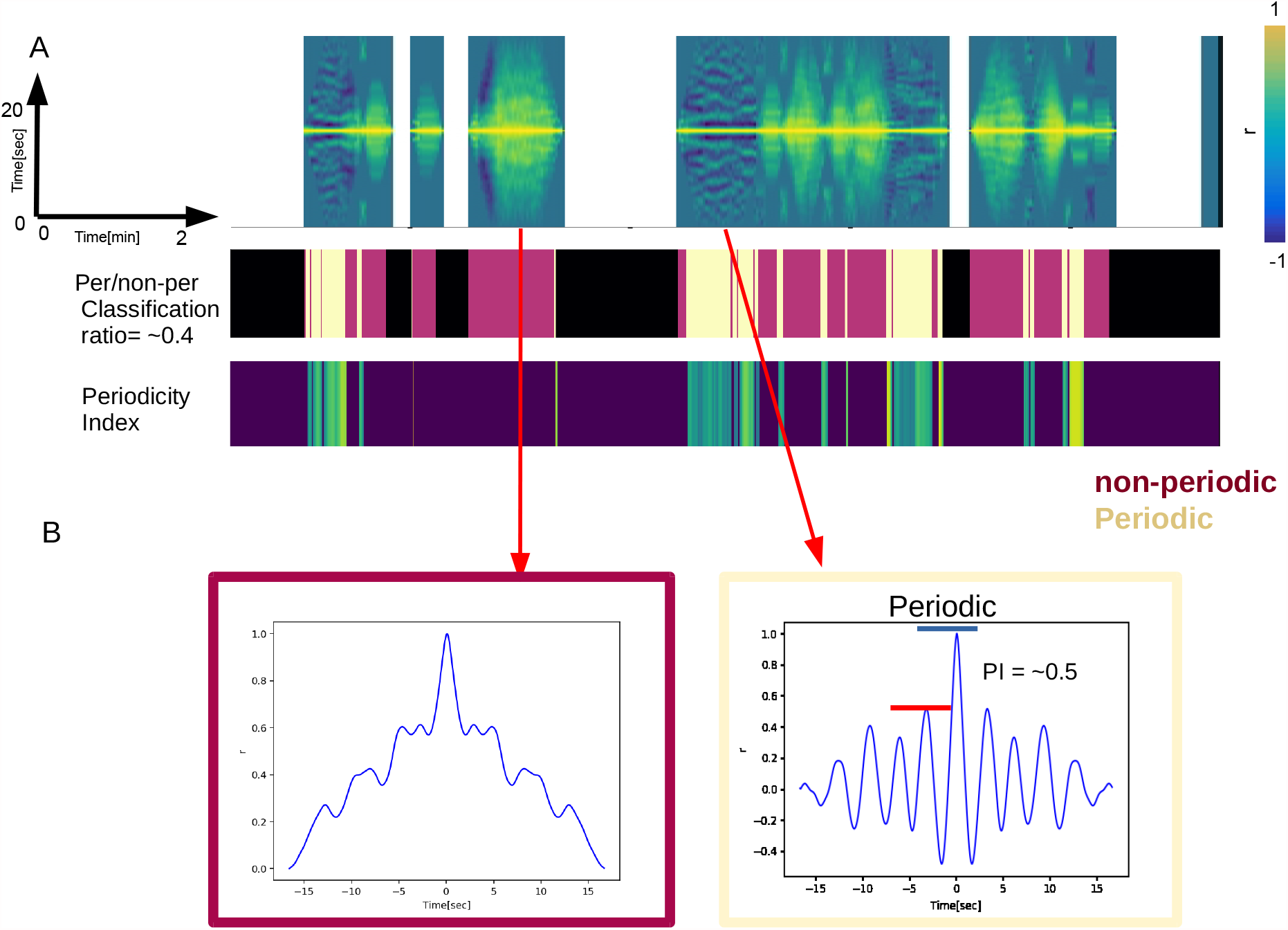
Periodicity Index is used to measure the strength of periodicity and to separate periodic from non-periodic behaviors. **(A)** Examples of autocorrelation functions (ACFs) of the long time-scale as they evolve over 20 minutes (top), the corresponding classification of periodic (Chardonnay color) and non-periodic behaviors (Pinot Noir color) (middle) and the Periodicity Indexes of the periodic behaviors (bottom). (**B**) Examples of ACFs computed from non-periodic (left) and periodic (right) long time-scale behaviors. The blue and red bars indicate the heights of the central peak and its nearest neighbor respectively. Periodicity Index is only defined for behaviors classified as periodic (right). A sine wave would have the Periodicity Index of 1. Less periodic behaviors have the PI approaching zero.

**Figure 1 – figure supplement 3:**
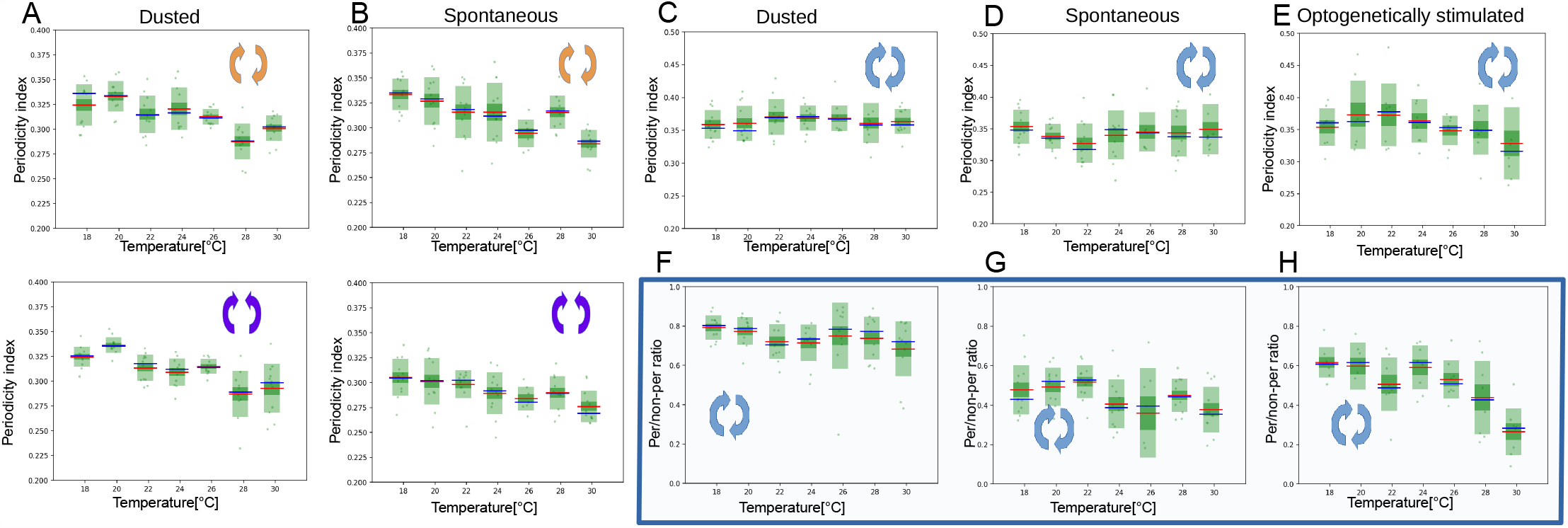
Analysis of periodicity of both time-scales for dust-stimulated flies, spontaneously grooming and optogenetically stimulated flies. (**A**) Periodicity indexes for the short time-scale of dust-stimulated flies across the seven temperatures (top: leg rubbing, bottom: head cleaning). (**B**) Similar as in (A) but for the spontaneously grooming flies. (**C**) to (**E**) Periodicity indexes for the long time-scale of dust-stimulated, spontaneously grooming flies and optogenetically-stimulated flies respectively. (**F**) to (**H**) Ratios of periodic to non-periodic long time-scale behaviors of dust-stimulated, spontaneously grooming and optogenetically-stimulated flies respectively.

**Figure 1—figure supplement 4:**
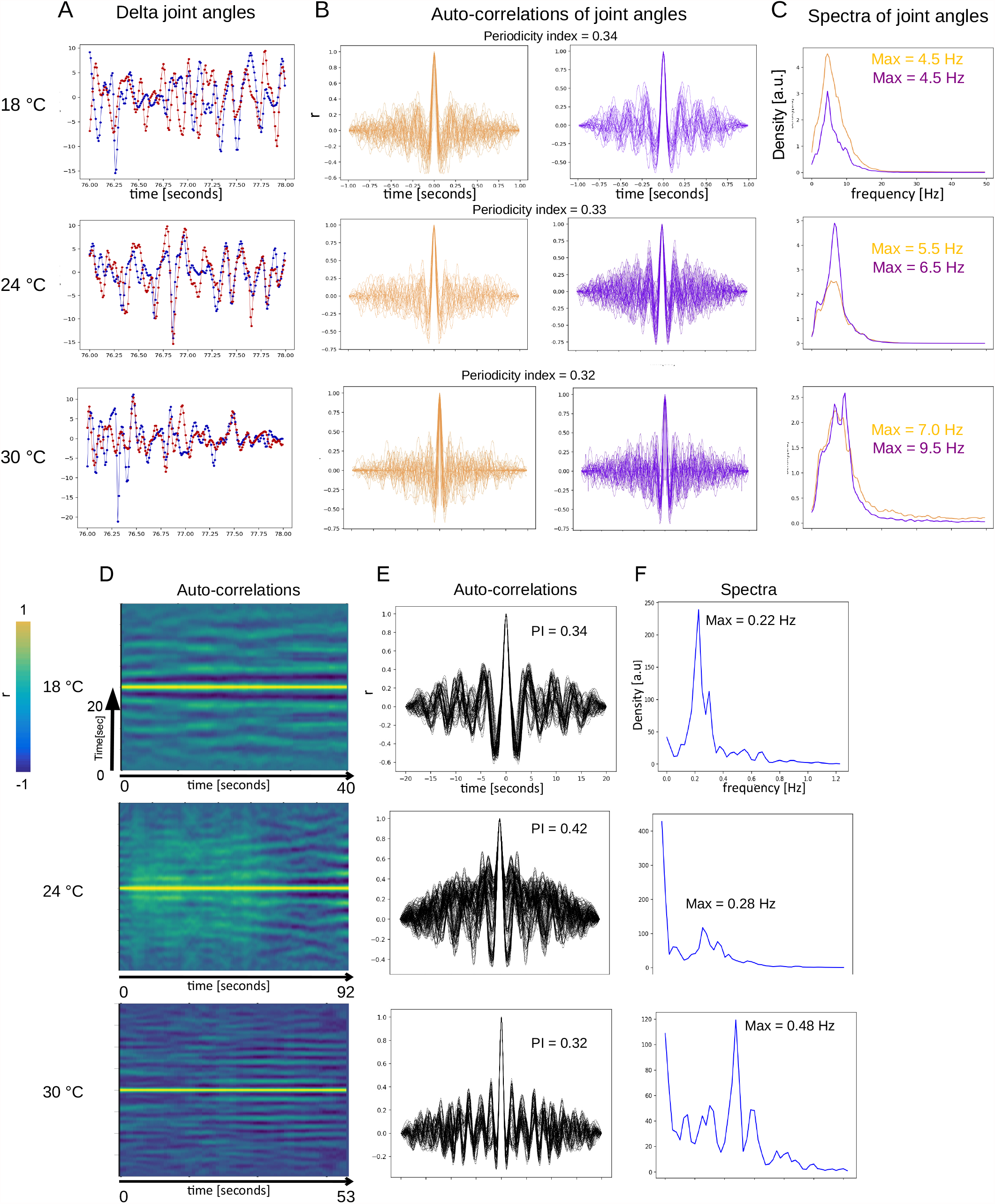
Periodicity of both time-scales is confirmed by an independent method of behavior analysis. (**A**) Examples of time-traces of angle changes of two joints from opposite front legs (red and blue traces) from flies recorded at three different temperatures (18 C, 24 C and 30 C). (**B**) Auto-correlation functions (ACFs) of the time-traces sampled from same three temperatures. Orange = presumptive front-leg rubbing, purple = presumptive head-sweeps. (**C**) Spectra of the same data as in (B). Flies in (A), (B) and (C) are optogenetically stimulated. (**D**) Auto-correlation functions of the long time-scale plotted across time of dust-stimulated flies at the three temperatures. The input for the auto-correlation analysis was a signal constructed from the joint positions as described in the Methods. (**E**) Same data as in (D) but shown as overlaid ACFs. Notice the increase of the number of side-peaks with temperature indicating increase of frequency. (**F**) Spectra of the long time-scale (same data as in (D) and (E)) showing the temperature-dependent max frequency increase.

**Figure 2 – figure supplement 1:**
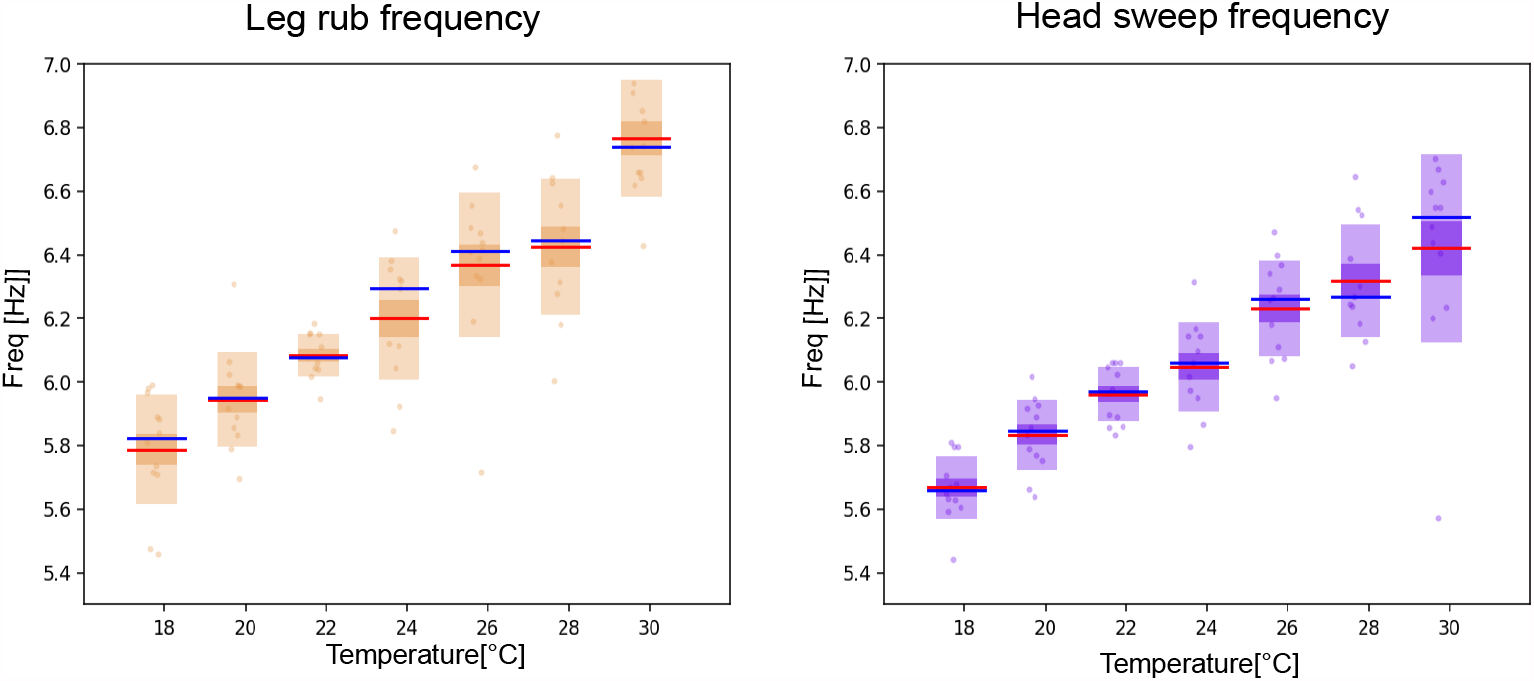
Frequencies of leg rubs and head sweeps increase with temperature. Separating the individual leg movements into leg rubs and head sweeps shows that the frequency of both increases with temperature at a similar rate.

**Figure 2 – figure supplement 2:**
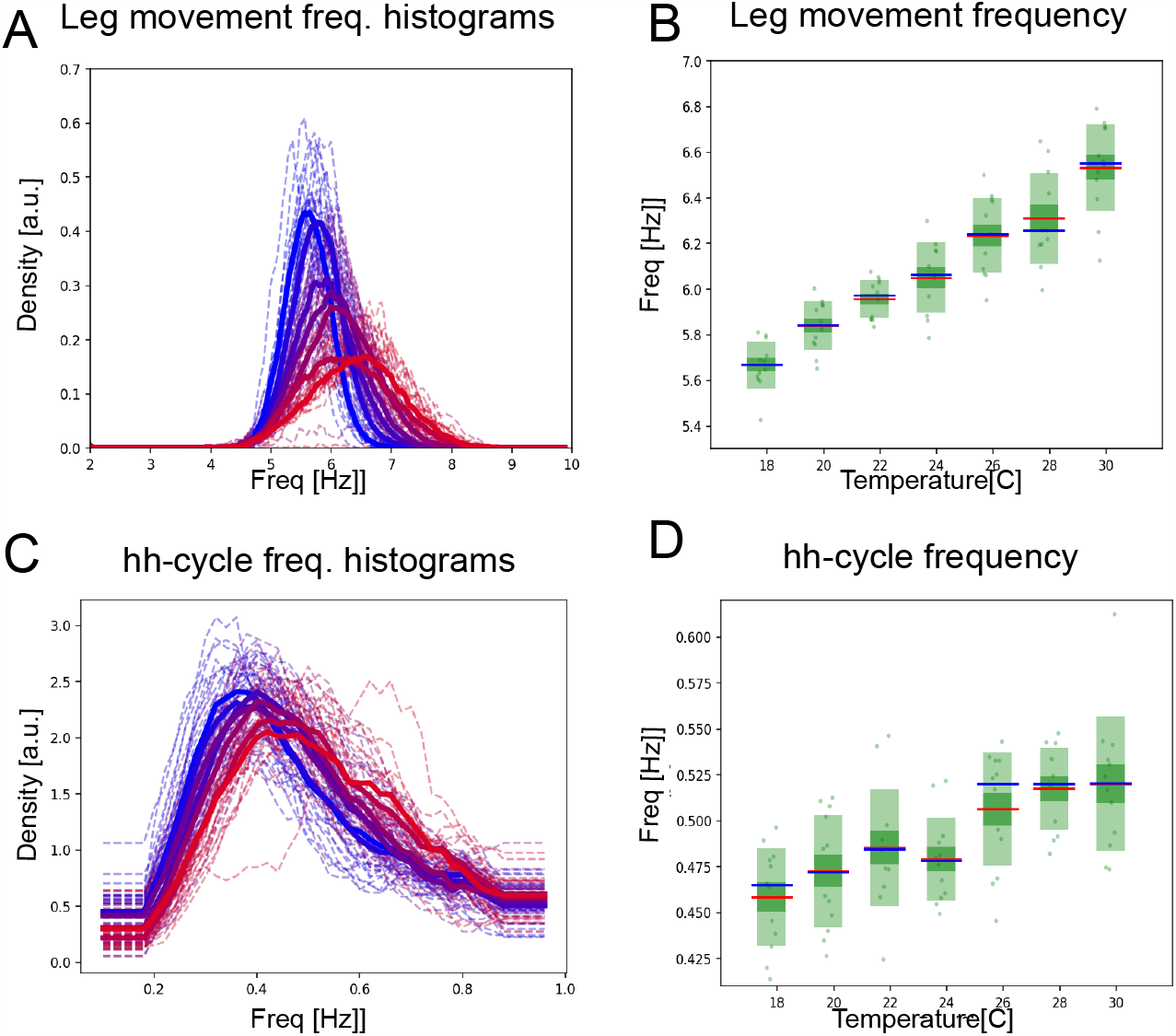
The hh-cycles also contract with increasing temperature. (**A**) Histograms of mean frequencies at cool (blue) and warm (red) temperatures, and (**B**) box plots of the increase in leg movement frequency with temperature; plots and statistics as described in **Figure 2 D**, and **E. (C)** Long-time scale hh-cycle analysis comparable to **Figure 2 G** and **H**.

**Figure 2 – figure supplement 3:**
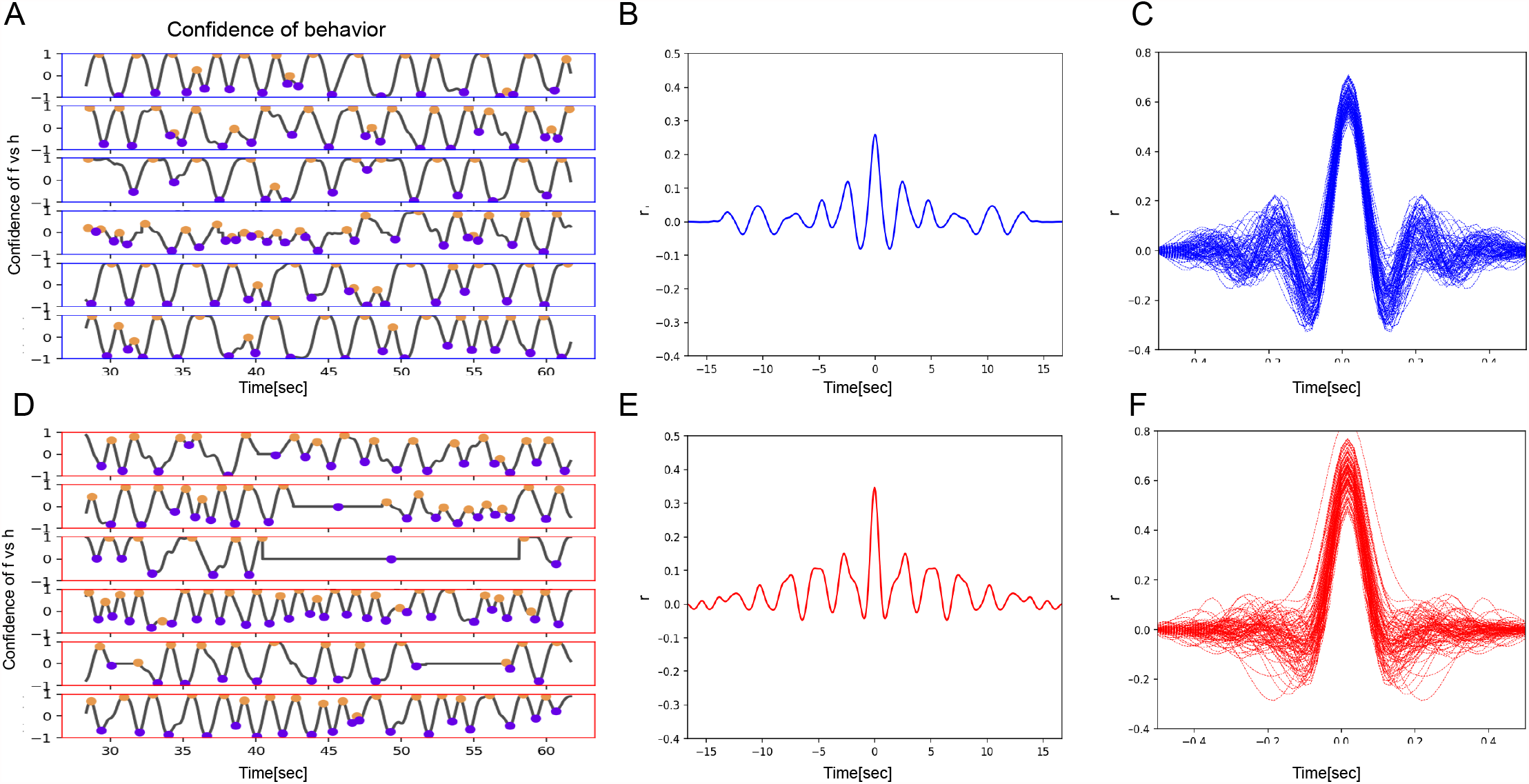
Bout alternations remain periodic across a range of temperatures. (A) Samples of five flies recorded at low temperature (18°C, blue frame), showing behavior confidence curves with several ff/hh cycles. Purple and orange circles indicate the times of h- and f-peaks. (**B**) Autocorrelation function curves of behavior confidence sampled from 18°C flies, indicating strong periodicity of the signals (side peaks). (**C**) Samples of autocorrelation functions (ACF) computed from over 8 minutes of a movie when the fly was engaged in front leg rubbing or head sweeps. (**D**)-(**F**) Same as A-C but for high temperature (30°C).

**Figure 2 – figure supplement 4:**
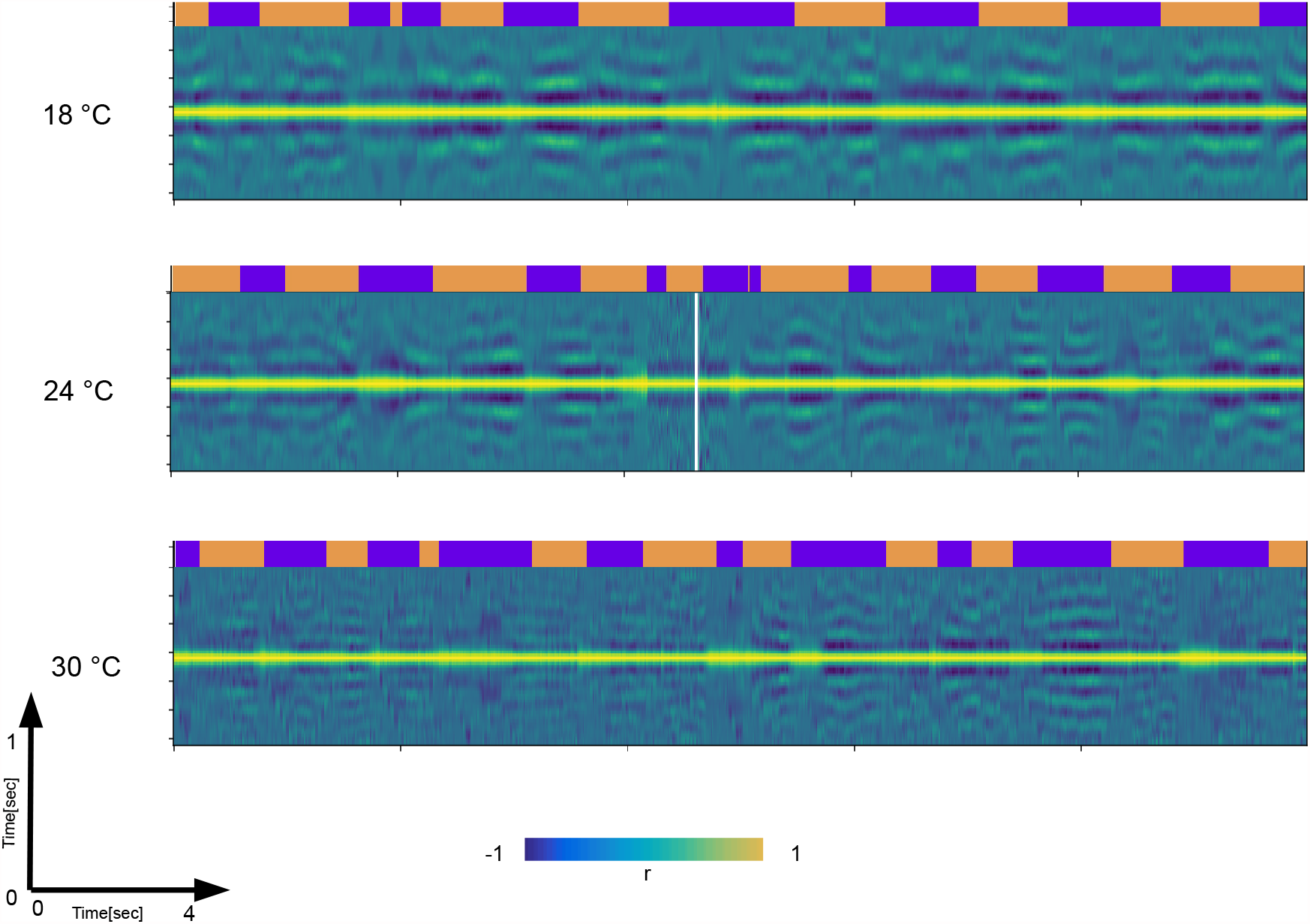
Autocorrelation analysis show temperature driven contraction across time-scales. Autocorrelation functions (ACFs) computed from 0.5 sec windows of raw videos of dusted flies are stacked together as columns into matrices across ~17 seconds of a movie. Each matrix is showing the change of ACF shape across the ~17 seconds. Note the modulation of the side-band positions and their numbers which is reflective of frequency modulation on the short time-scale (y-axis). Also note the modulation of the longer time-scales (x-axis). The three matrices shown are taken from different temperatures. Note that at the highest temperature the y-axis becomes “denser” (more side-bands) and that the episodes of harmonic bouts become shorter (x-axis). This way we can observe simultaneous transformation of x-axis and the y-axis as a result of temperature increase. The ethograms on top of the matrices are used as a reference.

**Figure 3 – figure supplement 1:**
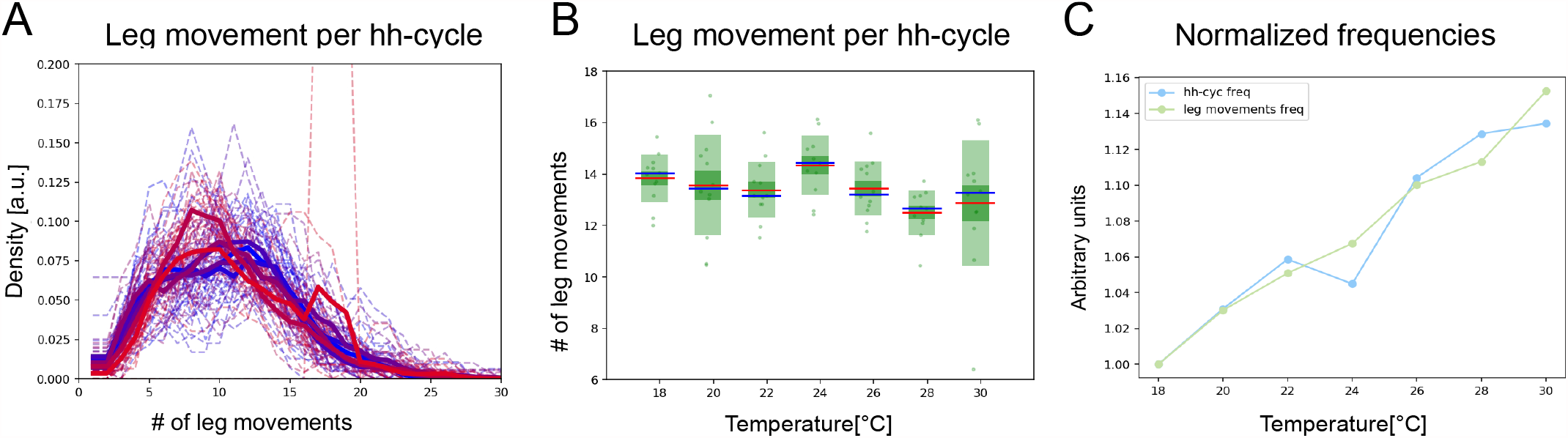
Head sweeps and hh-cycles contract at the same rate as temperature increases. **(A)** Histograms of numbers of leg movements per hh-cycle in cooler temperatures (blue) and warmer temperatures (red). **(B)** Box plots of leg movement counts per hh-cycle across the seven temperatures; statistics as in **Figure 2E**. (**C**) The frequency of individual leg movements and bout alternations (hh-cycles) increases roughly linearly with temperature but over different time-scales (msec vs. sec; 7Hz vs. 0.5Hz).

**Figure 4 – figure supplement 1:**
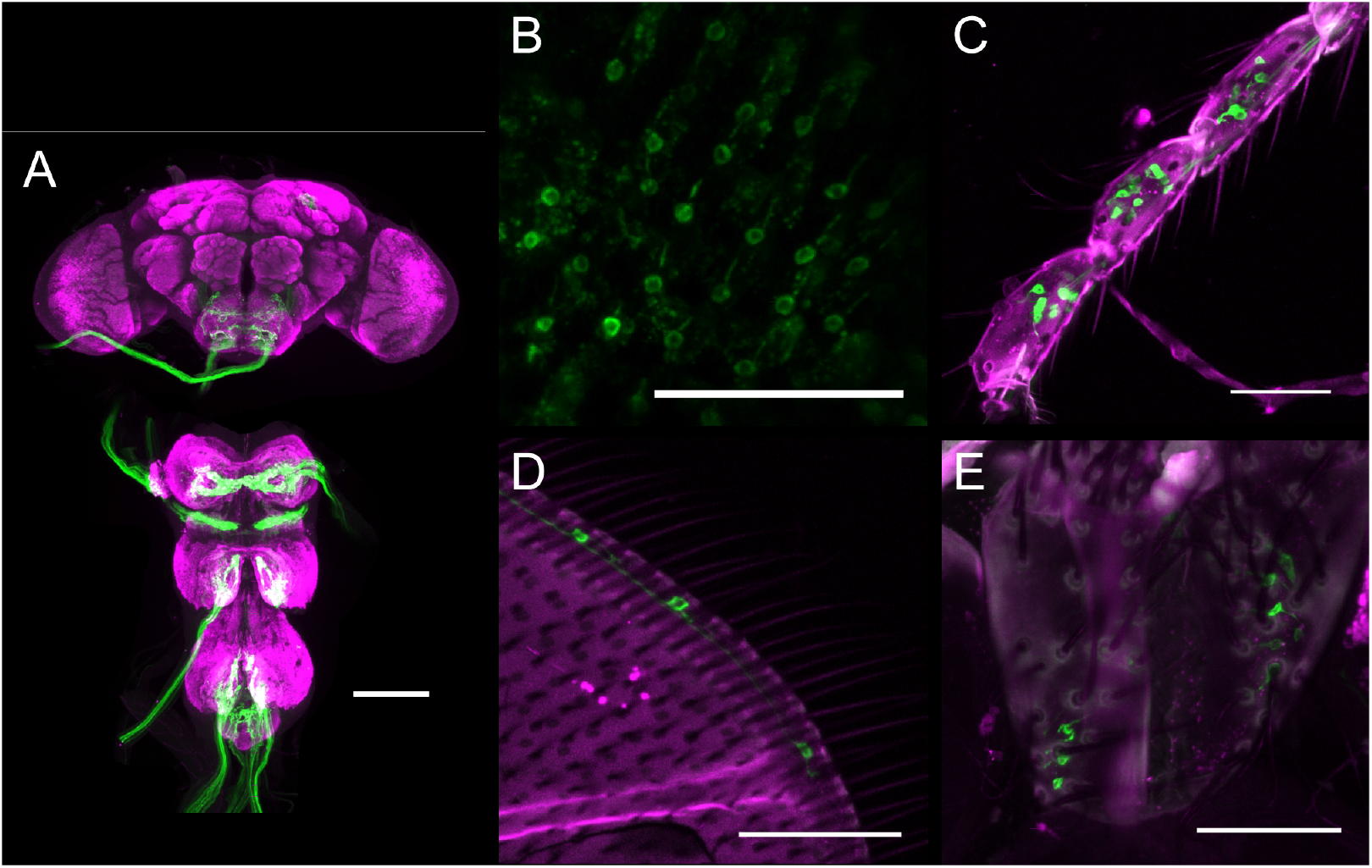
Expression pattern of the mechanosensory bristles driver line used in optogenetics experiments. (**A**) Expression pattern of R74C07-GAL4 in central nervous system. Magenta: anti-Bruchpilot. Green: anti-GFP. Scale bars, 100 mm. (**B**-**E**) Expression pattern of R74C07-GAL4 in eye bristles (B), leg bristles (C), wing bristles (D) and abdominal bristles (E). Magenta: cuticle autofluorescence. Green: innate GFP fluorescence. Scale bars, 50 mm.

**Figure 4 – figure supplement 2:**
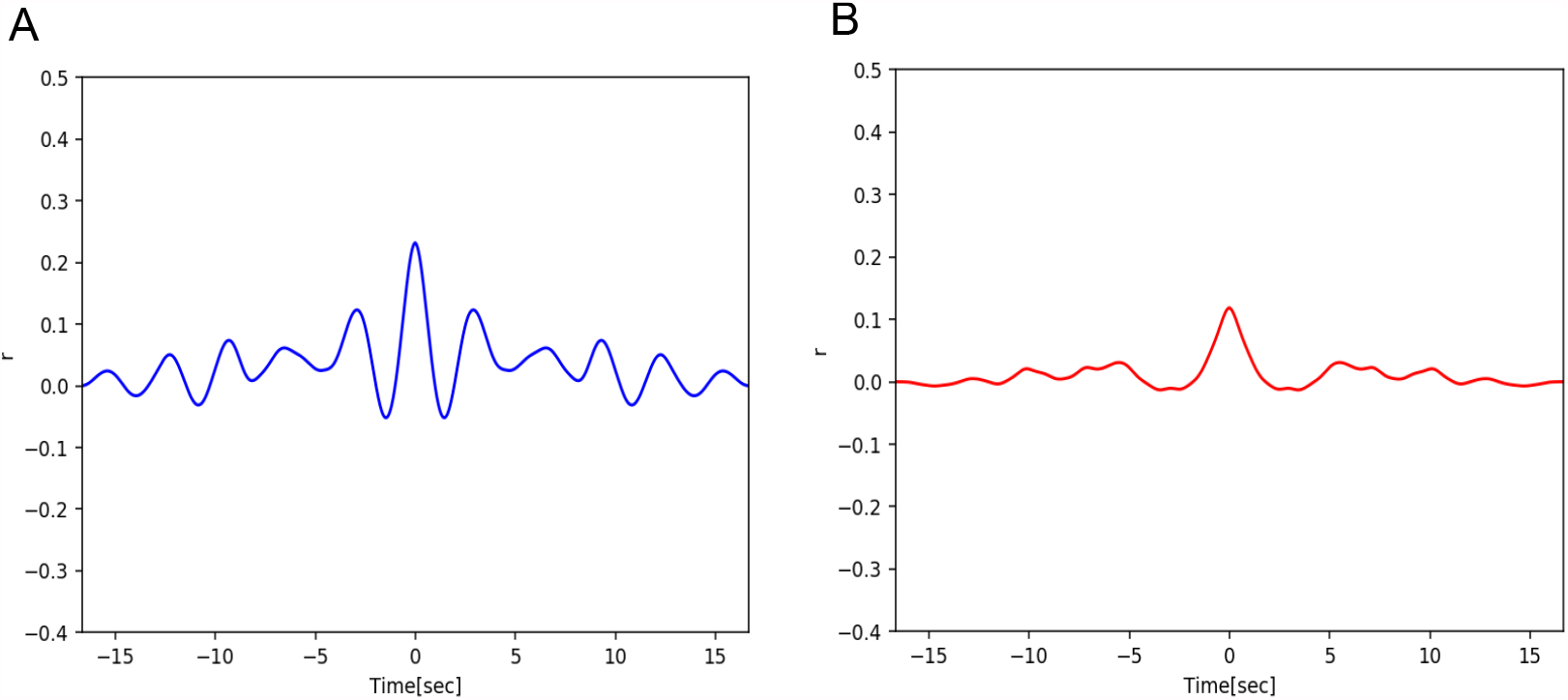
Bout alternations remain periodic across a range of temperatures in optogenetically-stimulated flies. **(A)** Autocorrelation function curves of behavior confidence sampled from 18°C flies, indicating strong periodicity of the signals (side peaks). (**B**) Same as **A** but for high temperature (30°C). The autocorrelations here are lower than in **A**, however side-peaks remain.

**Figure 4 – figure supplement 3:**
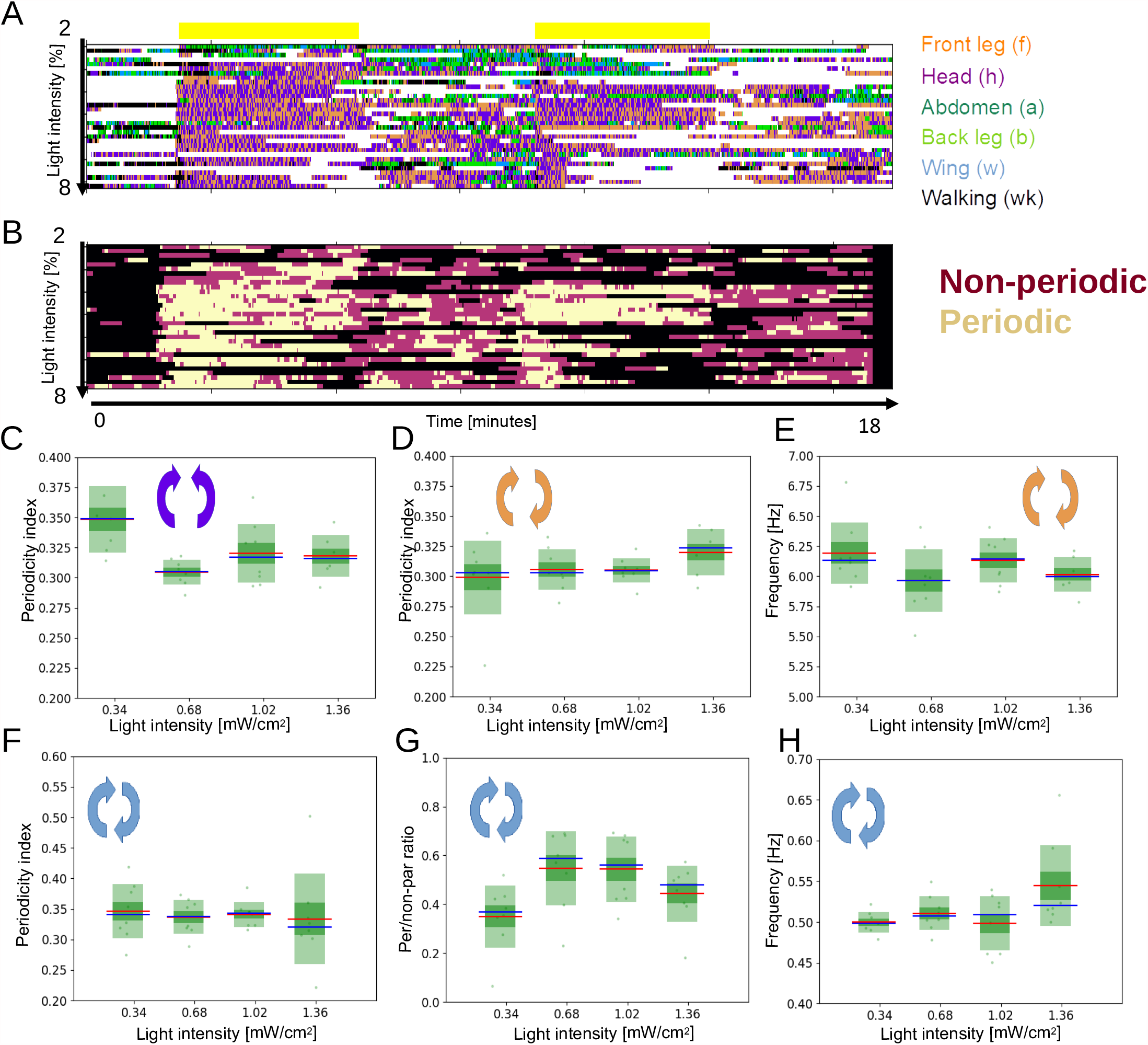
Optogenetically stimulated flies at different light intensities. (**A**) Ethograms representing 32 flies stimulated with different light intensities (0.34, 0.68, 1.02 and 1.36 mW/cm^2^). The yellow bars represent the periods of light activation, lasting 2 minutes each, to optogenetically induce grooming. (**B**) Ethograms showing periodic f-h alternations (beige color), non-periodic f-h alternations (purple) and other (non-anterior grooming) behaviors (black). (**C**) and (**D**) Periodicity indexes (PI) for head sweeps and leg rubs respectively, across the four light intensities. (**E**) Frequencies of leg rubs across the light intensity levels. (**F**) Periodicity Indexes for ff-cycles. (**G**) Periodic to non-periodic behavior ratio (for ff-cycles) across the four light intensities. (**H**) Similar as in **E** but for the long time-scale (ff-cycles).

**Figure 5 – figure supplement 1:**
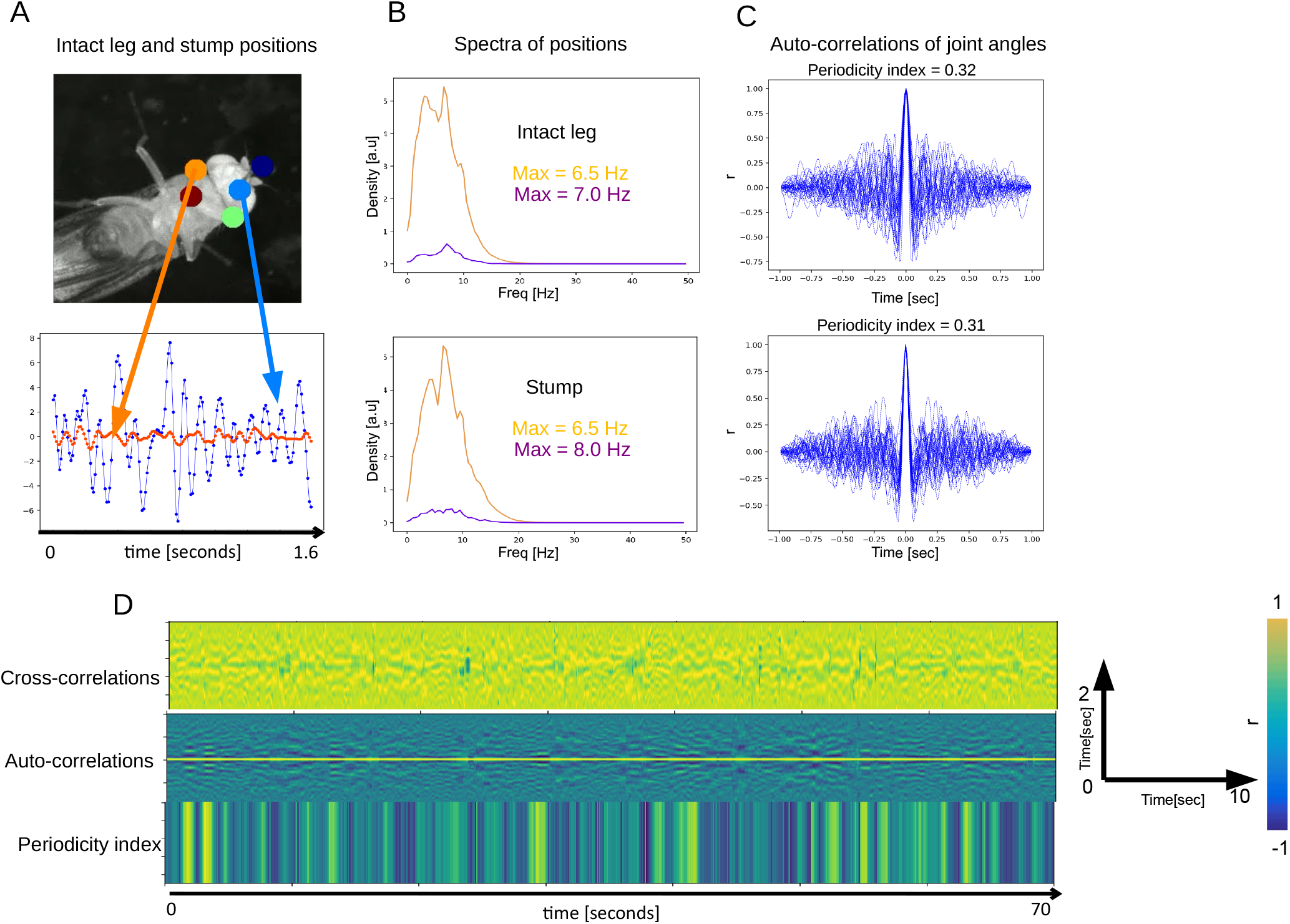
Periodicity and frequency is preserved in amputated front leg. (**A**) Top: positions of joints and the stump (orange) annotated by Deep Lab Cut (DLC). Bottom: positions of a joint on the intact front leg (blue) and stump (orange) as they change over time. (**B**) Spectra of position traces (over time) for the intact leg’s joint (top) and the stump (bottom). Gold and purple spectra correspond to the presumptive front-leg rubbing and head cleaning behaviors respectively. (**C**) Auto-correlation functions computed from the intact leg position traces (top) and from stump traces. (**D**) Cross-correlations between the intact leg and the stump (top), same auto-correlation functions as in **C** (middle) and the corresponding Periodicity Indexes (bottom) over 70 seconds.

**Figure 6 – figure supplement 1:**
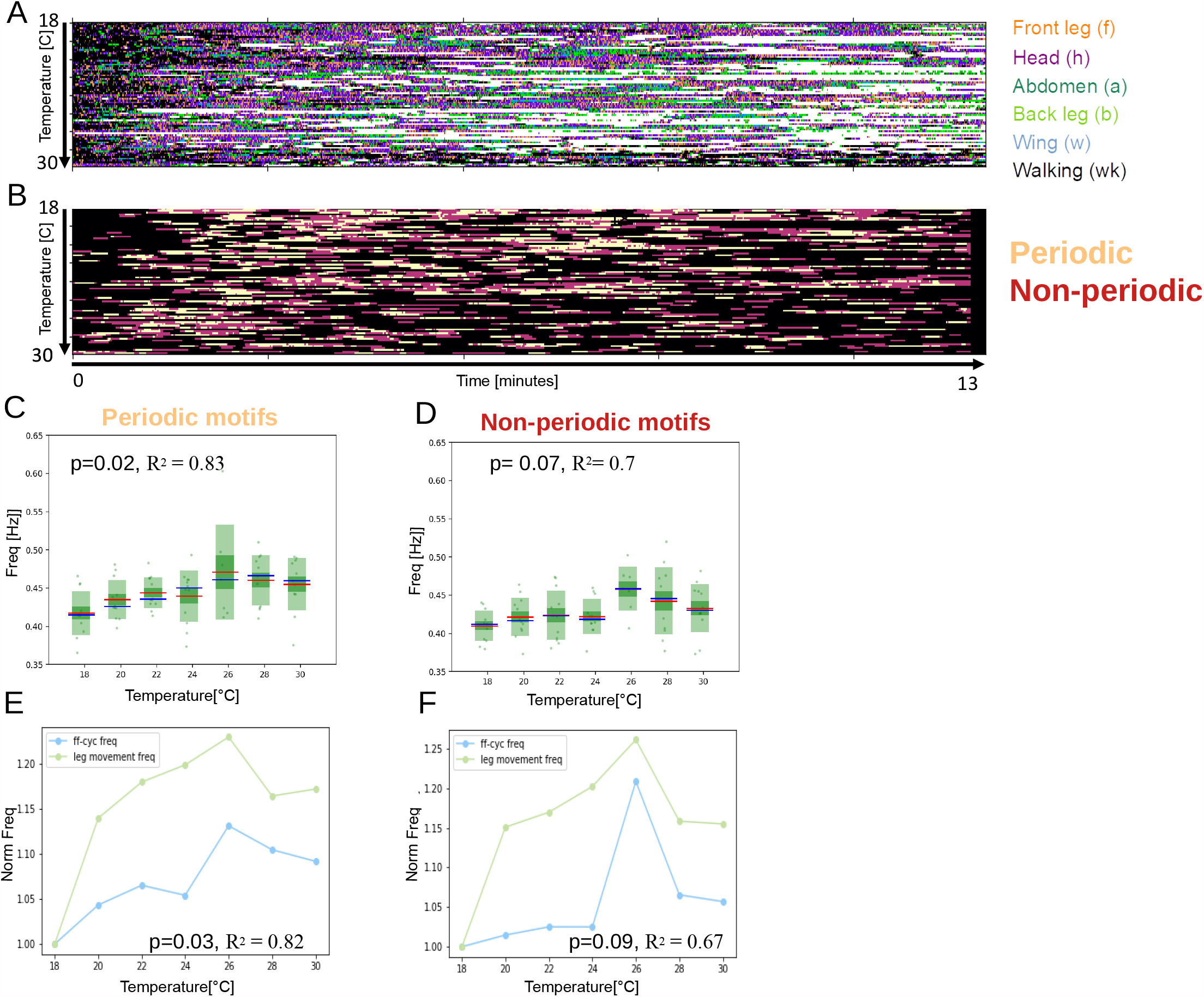
Periodic alternations between front leg and head cleaning scale with temperature better than the non-periodic behaviors in unstimulated flies. (**A**) Grooming and walking ethograms of unstimulated flies across the temperatures. (**B**) Ethograms showing periodic f-h alternations (beige color), non-periodic f-h alternations (purple) and other (non-anterior grooming) behaviors (black). (**C**) Box-plots of ff-cycle frequencies (as in **Figure 2E**) for the periodic f-h alternations (p=0.02, R^2^ = 0.83). (**D**) Same as **C** but for non-periodic f-h alternations (p= 0.07, R^2^= 0.7). (**E**) Normalized frequencies of leg movements and ff-cycle freq (similar as in **Figure 3C**) sampled from periodic grooming motifs. (**F**) Same as **E** but sampled from non-periodic grooming motifs.

**Figure 6 – figure supplement 2:**
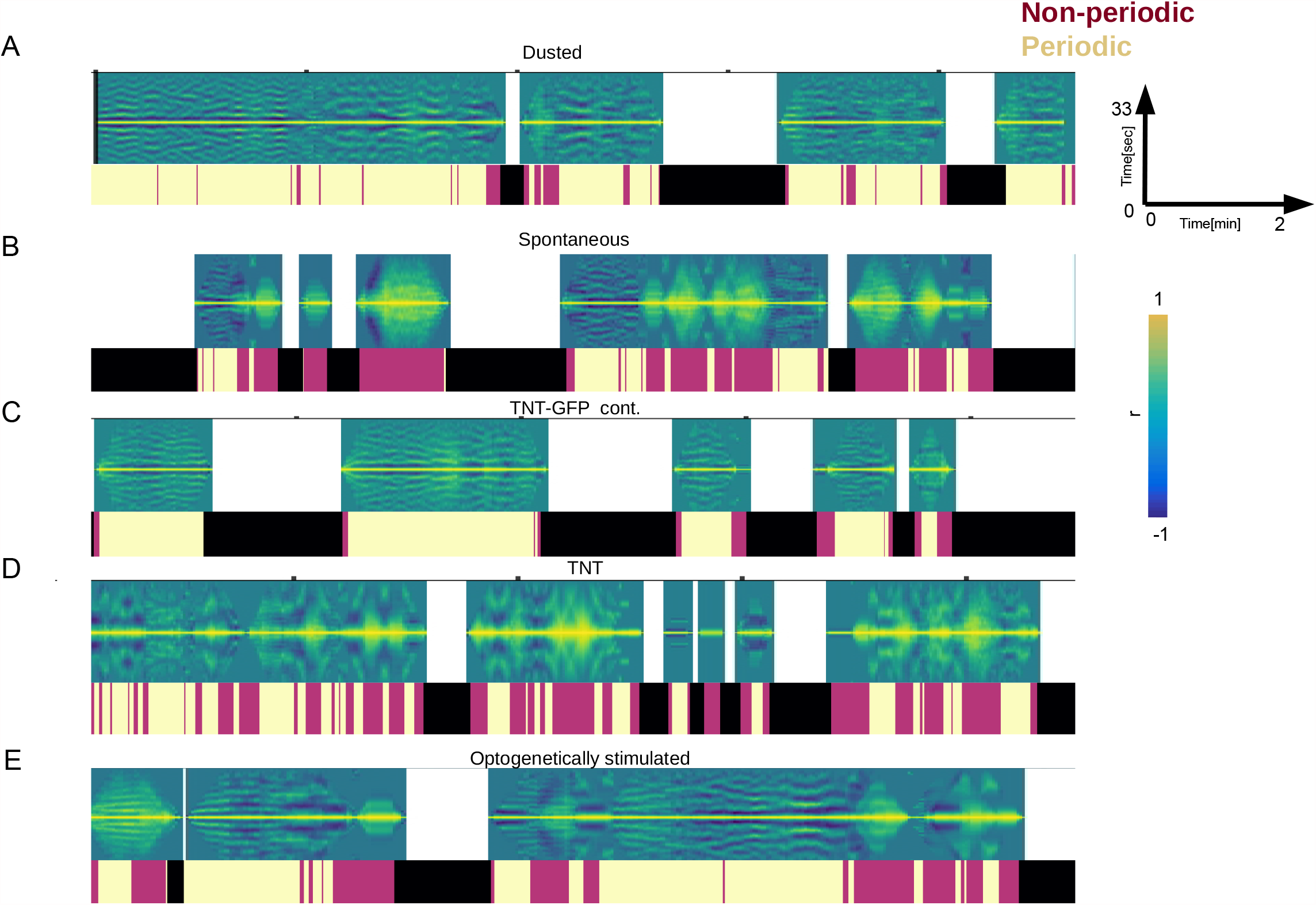
Periodicity of the long time-scale across different groups of flies. **(A)** An example of auto-correlation functions from a 13-minute video of a dusted fly at 18 °C. Ethograms placed underneath are showing periodic f-h alternations (Chardonnay color), non-periodic f-h alternations (Pinot Noir color) and other (non-anterior grooming) behaviors (black). (**B**), (**C**) and (**D**) Same as **A** but for a spontaneous grooming fly, a TNT-GFP control and TNT experimental fly (see **Figure 5**). The **C** and **D** were recorded at 24 °C.

## References

Alonso, L. M., & Marder, E. (2020). Temperature compensation in a small rhythmic circuit. Elife, 9. doi:10.7554/eLife.55470

Armstrong, E., & Abarbanel, H. D. (2016). Model of the songbird nucleus HVC as a network of central pattern generators. J Neurophysiol, 116(5), 2405–2419. doi:10.1152/jn.00438.2016

Berendes V, Zill SN, Büschges A, Bockemühl T. (2016). Speed-dependent interplay between local pattern-generating activity and sensory signals during walking in Drosophila. J Exp Biol. 2016 Dec 1;219(Pt 23):3781–3793.

Berkowitz, A. (2019). Expanding our horizons: central pattern generation in the context of complex activity sequences. J Exp Biol, 222(Pt 20). doi:10.1242/jeb.192054

Bidaye, S. S., Bockemuhl, T., & Buschges, A. (2018). Six-legged walking in insects: how CPGs, peripheral feedback, and descending signals generate coordinated and adaptive motor rhythms. J Neurophysiol, 119(2), 459–475. doi:10.1152/jn.00658.2017

Chen, C. L., Hermans, L., Viswanathan, M. C., Fortun, D., Aymanns, F., Unser, M., … Ramdya, P. (2018). Imaging neural activity in the ventral nerve cord of behaving adult Drosophila. Nat Commun, 9(1), 4390. doi:10.1038/s41467-018-06857-z

Deliagina, T. G., Orlovsky, G. N., & Pavlova, G. A. (1983). The capacity for generation of rhythmic oscillations is distributed in the lumbosacral spinal cord of the cat. Exp Brain Res, 53(1), 81–90. doi:10.1007/BF00239400

Feng, K., Sen, R., Minegishi, R., Dübbert, M., Bockemühl, T., Büschges, A., & Dickson, B. J. (2020). Distributed control of motor circuits for backward walking in <em>Drosophila</em>. bioRxiv, 2020.2007.2011.198663. doi:10.1101/2020.07.11.198663

Grillner, S. (2006). Biological pattern generation: the cellular and computational logic of networks in motion. Neuron, 52(5), 751–766. doi:10.1016/j.neuron.2006.11.008

Hampel, S., McKellar, C. E., Simpson, J. H., & Seeds, A. M. (2017). Simultaneous activation of parallel sensory pathways promotes a grooming sequence in Drosophila. Elife, 6. doi:10.7554/eLife.28804

Hughes, C. L. & J. B. Thomas. (2007) A sensory feedback circuit coordinates muscle activity in Drosophila. Molecular and Cellular Neuroscience 35.2: 383–396.

Jenett, A., Rubin, G. M., Ngo, T. T., Shepherd, D., Murphy, C., Dionne, H., … Zugates, C. T. (2012). A GAL4-driver line resource for Drosophila neurobiology. Cell Rep, 2(4), 991–1001. doi:10.1016/j.celrep.2012.09.011

Kaplan, H. S., Thula, O. S., Khoss, N., & Zimmer, M. (2020). Nested neuronal dynamics orchestrate a behavioral hierarchy across timescales. Neuron, 105(3), 562–576.

Kays, I., Cvetkovska, V., & Chen, B. E. (2014). Structural and functional analysis of single neurons to correlate synaptic connectivity with grooming behavior. Nat Protoc, 9(1), 1–10. doi:10.1038/nprot.2013.157

Kiehn, O. (2016). Decoding the organization of spinal circuits that control locomotion. Nat Rev Neurosci, 17(4), 224–238. doi:10.1038/nrn.2016.9

Klapoetke, N. C., Murata, Y., Kim, S. S., Pulver, S. R., Birdsey-Benson, A., Cho, Y. K., … Boyden, E. S. (2014). Independent optical excitation of distinct neural populations. Nat Methods, 11(3), 338– 346. doi:10.1038/nmeth.2836

Long, M. A., & Fee, M. S. (2008). Using temperature to analyse temporal dynamics in the songbird motor pathway. Nature, 456(7219), 189–194. doi:10.1038/nature07448

Mantziaris, C., Bockemuhl, T., & Buschges, A. (2020). Central pattern generating networks in insect locomotion. Dev Neurobiol, 80(1-2), 16–30. doi:10.1002/dneu.22738

Marder, E., & Calabrese, R. L. (1996). Principles of rhythmic motor pattern generation. Physiol Rev, 76(3), 687–717. doi:10.1152/physrev.1996.76.3.687

Mathis A, Mamidanna P, Cury KM, Abe T, Murthy VN, Mathis MW, Bethge M. DeepLabCut: markerless pose estimation of user-defined body parts with deep learning. (2018). Nat Neurosci. 2018 Sep;21(9):1281–1289.

Mayer, W. P., & Akay, T. (2018). Stumbling corrective reaction elicited by mechanical and electrical stimulation of the saphenous nerve in walking mice. The Journal of experimental biology, 221(Pt 13), jeb178095. https://doi.org/10.1242/jeb.178095

Mendes CS, Bartos I, Akay T, Márka S, Mann RS. (2013.) Quantification of gait parameters in freely walking wild type and sensory deprived Drosophila melanogaster. Elife. 2013 Jan 8;2:e00231.

Mueller, J. M., Ravbar, P., Simpson, J. H., & Carlson, J. M. (2019). Drosophila melanogaster grooming possesses syntax with distinct rules at different temporal scales. PLoS Comput Biol, 15(6), e1007105. doi:10.1371/journal.pcbi.1007105

Mulloney, B., & Smarandache, C. (2010). Fifty Years of CPGs: Two Neuroethological Papers that Shaped the Course of Neuroscience. Front Behav Neurosci, 4. doi:10.3389/fnbeh.2010.00045

Phelps JS, Hildebrand DGC, Graham BJ, Kuan AT, Thomas LA, Nguyen TM, Buhmann J, Azevedo AW, Sustar A, Agrawal S, Liu M, Shanny BL, Funke J, Tuthill JC, Lee WA. (2021). Reconstruction of motor control circuits in adult Drosophila using automated transmission electron microscopy. Cell, Feb 4;184(3):759–774.

Phillis, R. W., Bramlage, A. T., Wotus, C., Whittaker, A., Gramates, L. S., Seppala, D., … Murphey, R. K. (1993). Isolation of mutations affecting neural circuitry required for grooming behavior in Drosophila melanogaster. Genetics, 133(3), 581–592.

Pires, A., & Hoy, R. R. (1992a). Temperature coupling in cricket acoustic communication. I. Field and laboratory studies of temperature effects on calling song production and recognition in Gryllus firmus. J Comp Physiol A, 171(1), 69–78. doi:10.1007/BF00195962

Pires, A., & Hoy, R. R. (1992b). Temperature coupling in cricket acoustic communication. II. Localization of temperature effects on song production and recognition networks in Gryllus firmus. J Comp Physiol A, 171(1), 79–92. doi:10.1007/BF00195963

Powell, D. J., Haddad, S. A., Gorur-Shandilya, S., & Marder, E. (2020). Coupling between fast and slow oscillator circuits in Cancer borealis is temperature compensated. bioRxiv, 2020.2006.2026.173427. doi:10.1101/2020.06.26.173427

Pulver, S. R., Bayley, T. G., Taylor, A. L., Berni, J., Bate, M., & Hedwig, B. (2015). Imaging fictive locomotor patterns in larval Drosophila. J Neurophysiol, 114(5), 2564–2577. doi:10.1152/jn.00731.2015

Ravbar, P., Branson, K., & Simpson, J. H. (2019). An automatic behavior recognition system classifies animal behaviors using movements and their temporal context. J Neurosci Methods, 326, 108352. doi:10.1016/j.jneumeth.2019.108352

Rinberg, A., Taylor, A. L., & Marder, E. (2013). The effects of temperature on the stability of a neuronal oscillator. PLoS Comput Biol, 9(1), e1002857. doi:10.1371/journal.pcbi.1002857

Santuz, A., Akay, T., Mayer, W.P., Wells, T.L., Schroll, A. and Arampatzis, A. (2019), Modular organization of murine locomotor pattern in the presence and absence of sensory feedback from muscle spindles. J Physiol, 597: 3147–3165.

Sartoretti, G., Shaw, S., Lam, K., Fan, N., Travers, M., & Choset, H. (2018, May). Central pattern generator with inertial feedback for stable locomotion and climbing in unstructured terrain. In 2018 IEEE International Conference on Robotics and Automation (ICRA) (pp. 5769–5775). IEEE.

Seeds, A. M., Ravbar, P., Chung, P., Hampel, S., Midgley, F. M., Jr., Mensh, B. D., & Simpson, J. H. (2014). A suppression hierarchy among competing motor programs drives sequential grooming in Drosophila. Elife, 3, e02951. doi:10.7554/eLife.02951

Selverston, A. I. (2010). Invertebrate central pattern generator circuits. Philos Trans R Soc Lond B Biol Sci, 365(1551), 2329–2345. doi:10.1098/rstb.2009.0270

Simpson, J. H., & Looger, L. L. (2018). Functional Imaging and Optogenetics in Drosophila. Genetics, 208(4), 1291–1309. doi:10.1534/genetics.117.300228

Stockinger P, Kvitsiani D, Rotkopf S, Tirián L, Dickson BJ. (2005). Neural circuitry that governs Drosophila male courtship behavior. Cell. 2005 Jun 3;121(5):795–807.

Tang, L. S., Taylor, A. L., Rinberg, A., & Marder, E. (2012). Robustness of a rhythmic circuit to short- and long-term temperature changes. J Neurosci, 32(29), 10075–10085. doi:10.1523/JNEUROSCI.1443-12.2012

Vandervorst, P., & Ghysen, A. (1980). Genetic control of sensory connections in Drosophila. Nature, 286(5768), 65–67. doi:10.1038/286065a0

Wang, M., Yu, J., & Tan, M. (2014). CPG-based sensory feedback control for bio-inspired multimodal swimming. International Journal of Advanced Robotic Systems, 11(10), 170.

Yamaguchi, A., Gooler, D., Herrold, A., Patel, S., & Pong, W. W. (2008). Temperature-dependent regulation of vocal pattern generator. Journal of neurophysiology, 100(6), 3134–3143.

Yu, Z., & Thomas, P. J. (2021). Dynamical consequences of sensory feedback in a half-center oscillator coupled to a simple motor system. Biological Cybernetics, 115(2), 135–160.

Zhang, N., Guo, L., & Simpson, J. H. (2020). Spatial Comparisons of Mechanosensory Information Govern the Grooming Sequence in Drosophila. Curr Biol, 30(6), 988–1001 e1004. doi:10.1016/j.cub.2020.01.045

